# Antibody-Fcγ receptor-driven NOX2 activation during phagocytosis of tumor cells is suppressed by CD47, preventing CD8 T cell engagement and immunotherapeutic responses

**DOI:** 10.64898/2026.11.19.767930

**Authors:** Pratibha Sharma, Lauren J. Olson, Seuly Akter, Jadyn F. Leek, Jason G. Kerkvliet, Hallie M. Kueter, Esme Fowkes, Natalie W. Thiex, Stephen M. Thirdborough, Stephen A. Beers, Mark S. Cragg, Adam D. Hoppe

## Abstract

Antibody-dependent cellular phagocytosis (ADCP) contributes to the elimination of antibody-opsonized tumor cells and engagement of adaptive immunity. This process is inhibited by tumor-cell overexpression of CD47. Here, genome-wide CRISPR screens in macrophages identified the NADPH oxidase (NOX2 complex) and the CD47 receptor, SIRPα, as regulators of phagosome acidification. Target-cell CD47 suppressed NOX2 generation of reactive oxygen species at antibody densities that support ADCP, revealing distinct CD47-dependent thresholds for ADCP and oxidant generation. CD47 removal from the target cell delayed phagosome acidification in wild-type macrophages, but not in NOX2-deficient macrophages.

Consistent with a role for this pathway in antitumor immunity, tumor-opsonizing anti-Tyrp1 therapy allowed clearance of CD47-deficient B16F10 melanoma tumors in approximately 36% of wild-type mice but failed in NOX2-deficient mice. Tumor clearance required CD8 T cells and IFNγ, but perforin was dispensable. CD4 T cell depletion markedly improved tumor clearance. These findings suggest a noncanonical role for CD8 T cells in supporting ADCP-mediated tumor control.

Combining anti-Tyrp1 with a mouse-serum-albumin-IL-2 fusion (MSA-IL-2) or PD-1 blockade enabled superior clearance of CD47-deficient tumors in NOX2-deficient mice, but not wild-type mice. Rechallenge experiments distinguished primary tumor clearance from durable immunity, whereby NOX2-deficient mice treated with anti-Tyrp1 and PD-1 blockade lacked rechallenge protection despite high initial cure rates. Conversely, the addition of MSA-IL-2 supported durable protection in both wild-type and NOX2-deficient mice. These results indicate that tumor CD47 functions beyond a “don’t-eat-me” signal by controlling NOX2 activity to positively and negatively regulate CD 8 T cell function.

**Significance:** CD47-SIRPα suppresses NOX2 activation, regulating a critical feedback loop between ADCP and CD8 T cell activation during tumor-opsonizing antibody therapy, suggesting new strategies for cancer immunotherapy targeting tumor surface antigens.

## Introduction

Macrophages are abundant within solid tumors, where they can promote tumor progression and metastatic spread or, when appropriately activated, play a pivotal role in anti-tumor immune responses (Hussain et al. 2021), including improving the efficacy of T cell-based immunotherapies (Van Elsas et al. 2024). During antibody-dependent cellular phagocytosis (ADCP), macrophages and other Fcγ receptor (FcγR)-expressing phagocytes eliminate cells displaying IgG bound to surface antigens (Cao et al. 2022; Hirayama et al. 2017; Swanson and Hoppe 2004; Dooling et al. 2023). Therapeutic tumor-targeting antibodies have demonstrated efficacy against a variety of solid tumors and hematologic targets. For example, anti-CD20 antibodies drive phagocytic depletion of normal and malignant B cells (Marshall et al. 2017), while anti-HER2 antibodies support ADCP of HER2-positive breast cancer cells in addition to interfering with HER2 signaling (Tsao et al. 2019).

Macrophages recognize antibody-opsonized targets through FcγRs, which bind the Fc region of IgG and trigger phosphorylation of immunoreceptor tyrosine-based activation motifs (ITAMs), recruitment of Syk, and rearrangement of the actin cytoskeleton to engulf the target (Wöhner and Nimmerjahn 2025; Swanson and Hoppe 2004). In addition to initiating phagocytosis, FcγR engagement recruits and activates the phagocyte NADPH oxidase (NOX2), which produces reactive oxygen species (ROS) within the phagosome (Balce et al. 2016).

Following phagocytosis, antigens spared from degradation by lysosomal hydrolases can be presented on MHC molecules, supporting T cell activation. In the classic model, presentation of phagosome-derived antigens on MHC class II activates CD 4 T helper 1 cells that produce IFNγ, which in turn drives macrophage activation (Lau et al. 2025; Hodny et al. 2016). However, an emerging paradigm posits that cross-presentation of tumor antigens on MHC class I can support activation of CD8 T cells (Shanze Chen et al. 2023; Bode et al. 2023), which produce IFNγ, suggesting a parallel circuit. In turn, IFNγ-activated macrophages have increased phagocytic function and antigen presentation. Phagosomal antigens must be loaded onto MHC class I for cross-presentation and activation of CD8 T cells. NOX2-derived ROS preserves antigen and thus promotes cross-presentation by opposing phagosomal acidification through the consumption of H+ ions, inhibiting the vATPase, and permeabilizing the phagosome membrane (Savina et al. 2006; Mantegazza et al. 2008; Mpakali and Stratikos 2021). In addition, NOX2 oxidation of cysteines in cathepsins inhibits their protease function (Rybicka et al. 2010). NOX2 preservation of antigenic peptides facilitating their cross-presentation is best understood in classic dendritic cell type 1 (cDC1), which activate NOX2 downstream of lectin receptors such as DNGR-1 (Clec9) (Canton et al. 2021). FcγRs are known to activate NOX2 in the context of microbial infection; however, the presentation of cancer antigens following ADCP of tumor cells has not been carefully evaluated. Conversely, NOX2 activation in tumors may be a double-edged sword, as extracellular release of ROS causes oxidation of the T cell receptor and oxidative cell death of T cells (Gelderman et al. 2007; Belikov et al. 2015).

ADCP is antagonized by CD47, an integrin-associated cell surface protein expressed on nearly all cells of the body (Osorio et al. 2023; Bess et al. 2025). CD47 engages signal-regulatory protein alpha (SIRPα), which is selectively expressed at high levels on macrophages, monocytes, neutrophils, and classical dendritic cell type 2 (cDC2) cells (Willingham et al. 2012). Because many tumor types express high levels of CD47 (Kim et al. 2012; Majeti et al. 2009; Willingham et al. 2012), it has been intensely investigated as a therapeutically targetable immune checkpoint (Mirza et al. 2025; Willingham et al. 2012). However, systemic CD47 blockade causes toxicity, including anemia and splenomegaly, resulting from phagocytosis of red blood cells by macrophages (Mirza et al. 2025). Additionally, it has become increasingly clear that effective CD47 interventions require an opsonizing mechanism, such as an anti-tumor antibody or inclusion of chimeric-receptor antigen macrophages (Osorio et al. 2023). These complications have confounded its study as an immune regulator; however, the controlled production of tumor-localized CD47-blocking nanobodies demonstrates improved anti-tumor CD8 T cell activity (Siqi Chen et al. 2025), suggesting that CD47 signaling influences T cell engagement. Moreover, genetic removal of CD47 from tumor cells promotes anti-tumor antibody production during opsonizing immunotherapy (Dooling et al. 2023). Together, these findings argue that ADCP contributes to antigen processing and immune engagement, with CD47 serving as a critical checkpoint.

Here, we elucidated a new role for NOX2 in shaping tumor immunotherapy under the control of opsonizing antibodies and tumor-associated CD47. Specifically, we found that opsonizing antibodies overcome the target-cell CD47 suppression of ADCP for a wide range of targets and in solid tumors, particularly when macrophages are activated by IFNγ. However, even when IgG engagement of Fcγ receptors was sufficient to drive phagocytosis, CD47 on target cells strongly suppressed ROS production and accelerated phagosomal acidification, suggesting that CD47 imposes a higher threshold for antigen cross-presentation (a “don’t-present-me” signal) than for engulfment (a “don’t-eat-me” signal). Moreover, our findings suggest that during a strong immune reaction, anti-tumor IgG-driven NOX2 activation inhibits T cell responses during IL-2 augmentation or anti-PD-1 blockade immunotherapy.

## Methods

### Cell Culture

B16F10 (CRL-6475) and HEK293T (CRL-3216) cells were obtained from the American Type Culture Collection (ATCC). Cells were cultured in Dulbecco’s Modified Eagle’s Medium (DMEM; ATCC 30-2002) supplemented with 10% heat-inactivated fetal bovine serum (FBS; R&D Systems S11150H) and 1% penicillin-streptomycin (Pen-Strep; Gibco 15140-122) at 37℃ with 5% CO₂. Cells were detached using 0.05% trypsin-EDTA (Gibco 25300-054).

### Mice

C57BL/6 mice of various genotypes were used according to the experimental requirements. Wild - type (WT) C57BL/6 mice (Jackson Laboratory [JAX], stock no. 000664) served as controls. Other strains included NOX2-deficient (*Cybb*^-/-^) mice (B6.129S-*Cybb^tm1Din^*/J; JAX stock no. 002365), CD47-deficient (CD47^-/-^) mice (B6.129S7-*Cd47^tm1Fpl^*/J; JAX stock no. 003173), perforin1-deficient (*Prf1*^-/-^) mice (C57BL/6-*Prf1^tm1Sdz^*/J; JAX stock no. 002407), Cas9-transgenic mice (B6J.129(Cg)-*Gt(ROSA)26Sor^tm1.1(CAG-cas9*,-EGFP)Fezh^*/J; JAX stock no. 026179), and LyzM^CRE^ mice (B6.129P2-*Lyz2^tm1(cre)Ifo^*/J; JAX stock no. 004781) crossed with ZsGreen reporter mice (B6.Cg-*Gt(ROSA)26Sor^tm6(CAG-ZsGreen1)Hze^*/J; JAX stock no. 007906), resulting in LyzM^CRE^ x ZsGreen offspring. Mice were maintained in conventional, low-barrier individually ventilated cages. To allow environmental and physiological acclimation, mice were allowed at least 2 weeks of acclimation prior to experimental use. All animal experiments were conducted in accordance with protocols approved by the South Dakota State University Institutional Animal Care and Use Committee (IACUC). When possible, an equal number of male and female mice were used in experiments. No obvious sex differences were observed.

### Isolation and ex vivo culture of bone marrow-derived macrophages

Mice aged 8-14 weeks were euthanized by CO₂ inhalation followed by cervical dislocation. Bone marrow was collected from the femurs and tibias by flushing with Dulbecco’s phosphate-buffered saline (DPBS; Cytiva, SH30028.02). Bone marrow cells were cultured in a macrophage culture medium, consisting of DMEM supplemented with 20% FBS, 1% Pen-Strep, and 42 ng/mL macrophage colony-stimulating factor (M-CSF; BioLegend, 576406). Cells were incubated at 37°C in a humidified atmosphere with 5% CO₂. Additional media was added on days 3 and 5 and replaced on day 7, at which point cells had undergone differentiation into bone marrow - derived macrophages (BMDMs). For in vitro assays requiring inflammatory polarization, BMDMs were incubated with 50 ng/mL IFNγ (BioLegend, 575302) for 48 h, under the same conditions.

### Whole genome CRISPR screen

BMDMs were isolated from Cas9-transgenic mice and transduced on day 4 of differentiation with lentiviral particles containing the Brie mouse CRISPR knockout library (a gift from David Root and John Doench, Addgene, #73633-LV), which targets 19,674 genes with approximately four single-guide RNAs (sgRNAs) per gene and includes 1,000 non-targeting control sgRNAs (Doench et al. 2014). A total of 4 x 10⁷ BMDMs were transduced at a multiplicity of infection of 0.3 to favor integration of one or fewer sgRNA per cell. After 48 h, transduced cells were selected with puromycin (5 μg/mL) for 48 h, then cultured for an additional 2 days to allow depletion of the targeted proteins. Macrophages were polarized with IFNγ (50 ng/mL) for 48 h prior to the screen.

To assess phagosome acidification following ADCP, macrophages were plated in 10-cm non-tissue-culture-treated dishes (Fisher Scientific, FB0875712) at 2 x 10⁶ cells per dish. Macrophages were incubated with biotinylated mouse red blood cells (mRBCs) that were opsonized with 3 x 10^6^ anti-biotin IgG2a antibody (Abcam, ab36406) and labeled with pH-sensitive pHrodo Red-AM dye at a target-to-macrophage ratio of 30:1, then incubated at 37°C for 40 min. Following incubation, non-phagocytosed mRBCs were removed by washing with DPBS. Macrophages were detached using cold DPBS without calcium or magnesium, maintained on ice, and sorted by fluorescence-activated cell sorting using a BD FACSJazz cell sorter into populations with high and low pHrodo fluorescence, corresponding to greater and lower phagosome acidification, respectively.

### Genomic DNA extraction, Amplification, Sequencing and Analysis

Genomic DNA (gDNA) was extracted from sorted macrophage populations using the GeneJet Genomic DNA Isolation Kit (K0721), following the method previously described (Hossain et al. 2023). Integrated sgRNA sequences were amplified by PCR using Phire Green Hot Start II PCR Master Mix (Thermo Scientific, F126S). Each PCR reaction contained 25 µL of PCR master mix, 0.71 µL of 14 µM P5 stagger primer mix, 5 µL of 14 µM uniquely barcoded P7 primer mix, and 20 µL of gDNA solution. Thermal cycling consisted of an initial denaturation at 98°C for 30 sec, followed by 25 cycles of 98°C for 20 sec, 54°C for 15 sec, and 72°C for 20 sec, with a final extension at 72°C for 2 min. PCR products were pooled, separated by electrophoresis on a 2% agarose gel, and bands of the expected size were excised and purified using the Wizard SV Gel and PCR Clean-Up System (Promega, A9281). Purified amplicons were sequenced on an Illumina NextSeq 550 using a High Output 75-cycle sequencing kit. Sequencing reads were analyzed using the MAGeCK pipeline (Li et al. 2014) to quantify sgRNA abundance in each sorted population and identify sgRNAs enriched or depleted in high versus low pHrodo Red-fluorescence populations.

Differential pathway analysis was performed using the integrated Differential Expression and Pathway analysis (iDEP version 2.0, https://bioinformatics.sdstate.edu/idep20/). Data pre-processing and quality control were performed using default parameters, including missing value imputation via the gene median. Differentially expressed genes (DEGs) were identified using an FDR cutoff of 0.2 and a minimum fold-change threshold of two. To investigate functional implications, pathway enrichment analysis was performed on the identified gene sets using the Gene Ontology (GO) Molecular Function from the GSEA database. Pathway significance was evaluated using a false discovery rate (FDR) cutoff of 0.2 to 0.4. Enriched functional profiles, with a focus on downregulated genes, were visualized using lollipop plots generated within the platform, mapping pathway significance on the X-axis (-log10(FDR)), fold enrichment to node color, and gene counts to node sizes.

### Lentiviral transduction

Lentiviral packaging was performed in HEK293T cells (2 x 10⁶) seeded in a 10-cm tissue culture dish in DMEM supplemented with 10% FBS one day before transfection. The following day, cells were transfected with the lentiviral transfer, packaging, and envelope plasmids. For each transfection, 5 μg of the relevant transfer plasmid, either pLX_311-Cas9 (a gift from John Doench & William Hahn & David Root; Addgene plasmid #96924; http://n2t.net/addgene:96924 (Doench et al. 2014), LentiGuide-Puro encoding the CD47-targeting sgRNA (5’ - CCACATTACGGACGATGCAA - 3’) a gift from Feng Zhang (Addgene plasmid # 52963; http://n2t.net/addgene:52963), pLJM1-Lck-mScarlett (membrane localization of mScarlet; submission to Addgene pending) or pLenti nanoluciferase (NL; a gift from Erich Wanker; Addgene plasmid #113450; http://n2t.net/addgene:113450 (Trepte et al. 2018)), was combined with 3 μg of the packaging plasmid psPAX2 (a gift from Didier Trono; Addgene plasmid #12260; http://n2t.net/addgene:12260), 1 μg of the envelope plasmid pCMV-VSV-G (a gift from Bob Weinberg; Addgene plasmid #8454; http://n2t.net/addgene:8454 (Stewart et al. 2003)), and 24 μg of polyethyleneimine (PEI; PolySciences, catalog no. 23966). Following transfection, the cells were incubated at 37℃ and 5% CO₂. Lentiviral supernatants were harvested 48 h and 72 h after transfection and stored at −80°C until use. B16F10 cells were then exposed to lentiviral supernatants and after 24 h, the medium was replaced with DMEM supplemented with 10% FBS containing the appropriate selection antibiotic: 10 μg/mL blasticidin (Gibco, A11139-03) for Cas9-expressing cells, 2.5 µg/mL puromycin (Invivogen, cat. no. ant-pr) for cells transduced with CD47-targeting sgRNA, or 20 µg/mL Zeocin (Invitrogen, R25001) for nanoluciferase-expressing cells.

### Characterization of gene-edited cells

B16F10 CD47-knockout (B16F10^CD47-/-^) cells were generated as described above, harvested by trypsinization, and viable cells were counted by trypan blue exclusion. Cells were serially diluted in a 96-well plate to achieve limiting dilution and promote single-cell outgrowth. Plates were incubated at 37°C with 5% CO₂ and monitored periodically for the development of single-cell-derived colonies. Individual colonies were isolated and expanded sequentially in 24-well plates, 6-well plates, and 10-cm culture dishes. Clonal cell lines were subsequently assessed by flow cytometry. A total of 3 x 10⁵ cells were stained with anti-CD47 antibody (30 μg/mL; Bio X Cell, BE0270, clone MIAP301) in 100 μL of DPBS supplemented with 1% FBS for 30 min on ice. Cells were washed four times with DPBS containing 1% FBS, with centrifugation at 400 x g for 4 min per wash. Cells were then incubated with a 1:50 dilution of DyLight 488-conjugated goat anti-rat IgG secondary antibody (Invitrogen, SA5-10018) for 30 min on ice. Following three additional washes in DPBS, fluorescence was measured using the FL1/488 channel of a BD Accuri C6 flow cytometer.

Nanoluciferase expression in B16F10 and B16F10^CD47-/-^ cells was assessed by luminescence. Cells (2 x 10⁴) were seeded into an opaque white 96-well plate (Corning, 3789A) in DPBS, and the Vivazine substrate (Promega, N2580) was added at a 1:1,000 dilution. Luminescence was measured immediately with an Agilent BioTek Synergy H1 Hybrid Multi-Mode Reader.

### Cut Site Sequencing for CD47

Single-cell-cloned B16F10^CD47-/-^ cells were collected for genomic DNA analysis. Cell pellets containing 0.5 x 10⁶ to 2 x 10⁶ cells were collected by centrifugation, washed with DPBS, and processed for genomic DNA extraction using the GeneJET™ Genomic DNA Purification Kit (Thermo Scientific, K0721). The genomic region surrounding the CD47 CRISPR target site was amplified by PCR with primers designed using CHOPCHOP to generate an amplicon of approximately 350-450 bp. PCR was performed using Phire Hot Start II DNA Polymerase with the following primers: forward, 5′-TCCAGGTTCAGCTCAACTACTG-3′; reverse, 5′-TTCAAAACAGGCAAACTGTGTC-3′. PCR products were purified using AMPure XP magnetic beads (Beckman Coulter Life Sciences, A63881), quantified, and submitted to Plasmidsaurus for nanopore sequencing using its Premium PCR Sequencing service. Sequencing results were analyzed using CRISPResso2 (Clement et al. 2019) to characterize insertion and deletion (indel) frequencies at the target site.

### Luminol Assay

BMDMs were seeded at 15,000 cells per well in a white, opaque 96-well plate (Falcon, 0877126) in 200 μL of macrophage culture medium and allowed to adhere for 1.5 h at 37°C in a humidified atmosphere with 5% CO₂. Target cells, including mRBC, splenic B cells (SBC), or A20 cells, were opsonized with antibodies (anti-mouse 34-3C antibody (Invitrogen, MA5-49100) for mRBC, or anti-mouse CD20 antibody (clone 18B12; prepared in-house) for SBC and A20 cells) in Hanks’ balanced salt solution (HBSS; Cytiva, SH30268.01) supplemented with 1% FBS. A ROS detection solution containing 500 μM luminol (Sigma-Aldrich, A8511-5G), 50 U/mL horseradish peroxidase (HRP; Fisher Scientific, 51-653-15KU), and 14 mM dextrose in HBSS supplemented with 1% FBS was prepared. Opsonized target cells were mixed with the ROS detection solution at a 1:1 volume ratio, yielding final concentrations of 250 μM luminol, 25 U/mL HRP, and 7 mM dextrose. The culture medium was removed from the BMDMs, and 200 μL of the target cell (10 target cells per macrophage) and ROS detection mixture were added to each well. Luminescence was measured at 37°C at 5-min intervals for 1 h using an Agilent BioTek Synergy H1 Hybrid Multi-Mode Reader with luminescence fiber optics, a 3-sec integration time, and a read height of 1 mm.

### In vitro ADCP

BMDMs were differentiated for 7 days and, where indicated, polarized with IFNγ (50 ng/mL) for 48 h before the assay. A total of 30,000 macrophages were seeded per well in 96-well glass-bottom plates (Dot Scientific, MGB096-1-2-LG-L) and allowed to adhere for 2-3 h at 37°C in a humidified atmosphere with 5% CO₂. Target cells were labeled with pHrodo Red SE (Invitrogen, P36010) in DPBS without calcium or magnesium at concentrations optimized for each target cell type. Labeling was performed in a total volume of 250 μL for 40 min on a rotator. Following labeling, target cells (mRBC, SBC, or A20 cells) were opsonized with the appropriate antibody (anti-mouse 34-3C IgG2a antibody, Invitrogen, MA5-49100 for mRBC, or anti-mouse CD20 IgG2a antibody, clone 18B12 (prepared in-house) for SBC and A20 cells) at an optimized concentration in HBSS supplemented with 2% FBS for 40 min to facilitate FcγR-mediated recognition by macrophages. Labeled and opsonized target cells were then added to the macrophages (10 target cells per macrophage) using a multichannel pipette and incubated at 37°C. Incubation times were 30 min for smaller targets, such as mRBC, and 1 h for larger targets, such as SBC or tumor cells.

Fluorescence images were captured using an ImageXpress Micro XLS high-content imaging system (Molecular Devices) with a 20x air objective. Images were analyzed using a custom CellProfiler pipeline (Bray et al. 2015) to quantify target cell engulfment by macrophages. The phagocytic index (PI) was calculated as the measured area occupied by the target cells divided by the typical area of a single target cell, yielding an estimate of the number of target cells engulfed per macrophage.

### Phagosome Acidification

WT or *Cybb*^-/-^ BMDMs were differentiated for 7 days and, where indicated, polarized with 50 ng/mL IFNγ for 48 h before the assay. 20,000 macrophages were seeded per well in 96-well glass-bottom plates (Dot Scientific, MGB096-1-2-LG-L) and allowed to adhere for 2-3 h at 37°C in a humidified atmosphere containing 5% CO₂. 20-25 x 10⁶ mRBCs were labeled with 4 μM pHrodo Red SE (Invitrogen, P36010) and 16 μM Alexa Fluor 647 (AF647) NHS ester (Invitrogen, A37573) in DPBS without calcium or magnesium. Labeling was performed in a total volume of 500 μL for 30 min on a rotator. Labeled mRBCs were then opsonized with the 34-3C antibody (Invitrogen, MA5-49100) at a ratio of 10⁶ IgG molecules per RBC in HBSS supplemented with 1% FBS for 30 min. Calibration response curves were obtained by placing labeled mRBCs in PBS buffers of pH 4, 5.5, and 7.4 to determine pKa, Rmin, and Rmax. Fitting the response curve to pH = pKa -log10((Rmax-R)/(R-Rmin)) enabled image-based measurement of individual phagosomes. The opsonized mRBCs were added to the macrophages (5 target cells per macrophage) and incubated at 37°C for 30 min. Time-lapse fluorescence images were acquired over 6 h at 30 min intervals using an ImageXpress Micro XLS high-content imaging system (Molecular Devices) with a 20x air objective. Images were analyzed using a custom CellProfiler pipeline (Bray et al. 2015) to quantify phagosome acidification over time.

### Tumoroid Imaging

A 96-well V-bottom plate (Greiner Bio-One, 651161) was coated with anti-adherence rinsing solution (STEMCELL Technologies, 07010) for 20 min, washed with PBS, and blocked with DMEM supplemented with 10% FBS for 1 h. The blocking medium was removed, and 1,000 mScarlet-expressing B16F10^CD47-/-^ cells were seeded in 100 μL of complete DMEM per well on day 0 and incubated overnight at 37°C in a humidified atmosphere containing 5% CO₂. On day 1, 9,000 LyzM^CRE^ x ZsGreen BMDMs in 50 μL of macrophage medium were added to each well, with or without 2 μg of anti-Tyrp1 IgG2a antibody (Bio X Cell, BE0151, clone TA99). After overnight incubation, tumoroids were carefully transferred to a 96-well glass-bottom plate using a wide-bore 1-mL pipette and imaged on a Leica Stellaris 5 confocal microscope with 10x and 20x objectives.

### Luciferase Assay with tumoroids

A 96-well V-bottom plate (Greiner Bio-One, 651161) was coated with anti-adherence rinsing solution (STEMCELL Technologies, 07010) for 20 min, washed with DPBS, and blocked with DMEM supplemented with 10% FBS for 1 h. The blocking medium was removed, and 1,000 nanoluciferase-expressing B16F10^WT^ or B16F10^CD47-/-^ cells were seeded in 100 μL of culture medium per well on day 0 and incubated overnight at 37°C in a humidified atmosphere with 5% CO₂. On day 1, 9,000 BMDMs in 50 μL of macrophage culture medium were added to each well, with or without 2 μg of anti-Tyrp1 antibody (TA99). On day 3, the tumoroids were transferred to a solid-white 96-well plate (Corning, 3789A). Vivazine substrate (Promega, N2580) was added at 1:1,000 dilution, and the tumoroids were dissociated by pipetting. Luminescence was measured at 37°C using an Agilent BioTek Synergy H1 Hybrid Multi-Mode Reader, with kinetic readings acquired every 10 min for 3 h. Measurements used luminescence fiber optics, a 5-sec integration time, a gain of 150, and a read height of 1.5 mm. Luminescence at 110 min was used for subsequent analysis.

### Anti-CD20 B cell depletion

Mice (n = 3 per group) were bled on day 0, immediately before treatment. On days 1 and 5, mice received 50 μg of anti-CD20 antibody (clone 18B12, prepared in-house) in 100 μL of PBS containing calcium and magnesium via intravenous tail vein injection. In addition to anti-CD20 treatment, mice assigned to the CD47 blockade group received 500 μg of anti-CD47 antibody (Bio X Cell, BE0270; clone MIAP301) on days 0, 1, and 5. Blood samples (10-15 μL) were collected from the tail vein on days 3 and 7 using heparinized hematocrit tubes (Medline, CSX40B501). On day 11, mice were euthanized by CO₂ inhalation followed by cervical dislocation, and whole blood, spleens, and inguinal lymph nodes were collected for downstream analysis.

For flow cytometric assessment of circulating B cells, 10-15 μL of blood was diluted with 150 μL of DPBS supplemented with 1% FBS. Cells were stained with anti-CD45.2 APC (BioLegend, 109814) and anti-CD19 PE (BioLegend, 152408) at a 1:50 dilution in DPBS supplemented with 1% FBS and incubated on ice for 30 min. Red blood cells were lysed by adding 2 mL of ACK lysis buffer (Gibco, A1049201) and incubating at room temperature for 6 min and 30 sec. White blood cells were collected by centrifugation at 400 x g for 5 min at room temperature, washed with 1 mL of DPBS supplemented with 1% FBS, and centrifuged again at 400 x g for 5 min at 2-8°C. The resulting cell pellets were resuspended in DPBS for immediate flow cytometric analysis.

Spleens were mechanically dissociated in 500 μL of DPBS supplemented with 1% FBS and passed through a 30 μm cell strainer to obtain single-cell suspensions. An aliquot of 100 μL from each suspension was transferred to a fresh tube and stained with anti-CD45.2 APC (BioLegend, 109814) and anti-CD19 PE (BioLegend, 152408), each at a 1:50 dilution, then incubated on ice in the dark for 30 min. Red blood cells were lysed with 2 mL of ACK lysis buffer for 7 min at room temperature. Samples were centrifuged at 400 x g for 5 min, washed with 500 μL of DPBS, and centrifuged again at 400 x g for 5 min. Cell pellets were resuspended in 400 μL of DPBS and immediately analyzed by flow cytometry.

Inguinal lymph nodes were collected in 200 μL of DPBS supplemented with 1% FBS and mechanically dissociated with a 21-gauge needle or micro-pestle. The resulting suspensions were filtered through a 30 μm cell strainer. An aliquot of 100 μL from each suspension was transferred to a fresh tube and stained with anti-CD45.2 APC (BioLegend, 109814) and anti-CD19 PE (BioLegend, 152408) for 30 min on ice, protected from light. Samples were centrifuged at 400 x g for 5 min; the supernatant was removed, and cell pellets were resuspended in 400 μL of DPBS supplemented with 1% FBS for immediate flow cytometric analysis.

### In vivo tumor models

Mice aged 13-24 weeks were utilized for tumor challenges. The right flank of each mouse was shaved 1 day before tumor injection. B16F10^WT^ or B16F10^CD47-/-^ cells (2 x 10⁵) were injected subcutaneously into the right flank in 100 μL of HBSS with calcium and magnesium (Cytiva, SH30268.01). For anti-Tyrp1 monotherapy, mice received 250 μg of anti-Tyrp1 antibody (Bio X Cell, BE0151, clone TA99) intravenously in 100 μL of DPBS with calcium and magnesium on days 4, 5, 7, 9, 11, 13, and 15 after tumor injection. Control mice received mouse IgG2a isotype control antibody (Bio X Cell, BE0085) intravenously at the same dose and on the same schedule. For IFNγ-neutralization experiments, mice received 200 μg of anti-IFNγ antibody (Bio X Cell, BE0055, clone XMG1.2) intravenously on day 4, followed by 100 μg on days 5, 7, 9, 11, 13, and 15, in combination with anti-Tyrp1 antibody administered according to the schedule described above. For CD4 or CD8 T cell depletion experiments, mice received 300 μg of anti-CD4 antibody (clone GK1.5, prepared in-house) or 50 μg of anti-CD8 antibody (clone YTS169, prepared in-house) intravenously on days 1, 5, 11, and 17, in addition to anti-Tyrp1 antibody administered according to its standard schedule. For MSA-IL-2 treatment experiments, mice received 15 μg of MSA-IL-2 intravenously on days 4, 5, 7, and 13, either alone or in combination with anti-Tyrp1 treatment. For PD-1 blockade experiments, mice received 200 μg of anti-PD-1_Fcnull_ antibody (LALAPG; Syd Labs, PA007163.m2cLA, clone 29F.1A12.1) intravenously on days 4, 7, 11, and 15, either alone or in combination with anti-Tyrp1 treatment. Mice that survived the initial tumor challenge were rechallenged on day 80 with the same number of tumor cells without further treatment and monitored for tumor growth through day 160. Once palpable, tumors were measured using calipers every other day, and tumor area was calculated as (A = L x B x π/4), where (L) and (B) represent the longer and shorter tumor dimensions, respectively. Mice were euthanized by CO₂ inhalation followed by cervical dislocation when the tumor area approached the endpoint size of 125 mm².

### Plasma antibody binding assay

Blood was collected from experimental mice by terminal cardiac puncture or tail-vein sampling into heparinized hematocrit tubes (Medline, CSX40B501) and centrifuged at 800 x g for 5 min to obtain plasma. Plasma was collected and stored at -20°C until analysis. B16F10^CD47-/-^ cells (1.5 x 10⁵ per sample) were incubated with either 2 μg of anti-Tyrp1 antibody as a positive control or 2 μL of plasma from the respective mice in 100 μL of DPBS supplemented with 1% FBS for 30 min on ice. After staining, cells were washed four times with the same buffer, centrifuging at 400 x g for 4 min after each wash. Cells were then incubated with AF647-conjugated goat anti-mouse IgG secondary antibody (Invitrogen, A21236), diluted 1:200, for 30 min on ice. Following three washes with DPBS, cells were fixed in 4% paraformaldehyde and stored at 4°C until analysis. Fluorescence was measured using the FL4/APC channel on a BD Accuri C6 flow cytometer.

### MSA-IL-2 Protein Expression and Purification

Expi293F cells (Gibco, A14527) were cultured in Expi293 Expression Medium (Gibco, A1435101) at 37°C in a humidified atmosphere containing 8% CO₂. Cells were subcultured by centrifugation at 400 x g for 4 min and maintained at densities between 0.3 x 10⁶ and 3 x 10⁶ cells/mL. The MSA-IL-2 expression plasmid, a generous gift from Dr. K. Dane Wittrup (Zhu et al. 2015), was transfected into Expi293F cells using the ExpiFectamine Transfection Kit (Gibco, A14524) and Opti-MEM I Reduced Serum Medium (Gibco, 31985070), according to the manufacturer’s Expi293 Expression System protocol. Following expression for 5-7 days, the culture was centrifuged at 3,700 x g for 15 min to separate the cell pellet from the supernatant containing the secreted protein.

MSA-IL-2 was purified from the culture supernatant by immobilized metal affinity chromatography using His-tag Ni-NTA resin (Thermo Scientific, 88221). The protein was eluted with 0.25 M imidazole, desalted into phosphate-buffered saline (PBS), and further purified by size-exclusion chromatography using a HiPrep 16/60 Sephacryl S-200 HR column (Cytiva, 29497630) on an ÄKTA pure FPLC system. Protein concentration was determined using a bicinchoninic acid (BCA) assay (Thermo Scientific, 23235). Eluted fractions were analyzed by SDS-PAGE to assess protein size and purity, and protein ID was verified by trypsin digest and mass spectrometry. Endotoxin levels were measured using a chromogenic endotoxin quantification kit (Thermo Scientific, A39552), and preparations were required to meet an endotoxin limit of 0.3 EU per administered dose.

### Statistical Analysis

Statistical analyses were performed using GraphPad Prism (version 10; GraphPad Software, San Diego, CA) as indicated. Flow-cytometry data were analyzed using FlowJo (version 10.8.1; BD Biosciences, Franklin Lakes, NJ). The statistical tests used for individual experiments, along with the corresponding sample sizes, are indicated in the Figure legends. A *P* value of less than 0.05 was considered statistically significant.

## Results

### CRISPR whole-genome screens identified NOX2 regulation by the CD47 receptor, SIRPα

To elucidate how IFNγ supports macrophage ADCP, we conducted whole-genome CRISPR gene disruption screens. Cas9-transgenic BMDMs were mutagenized with the lentiviral sgRNA Brie library (Doench et al. 2014) and flow sorted based on the ability of resting (M0) or IFNγ-polarized macrophages to form acidic phagosomes following phagocytosis of IgG-opsonized mRBCs labeled with pH-sensitive pHrodo Red-AM (Fig. 1A). Deep-sequencing of the lentiviral sgRNA insertions was used to compare the frequency of mutant macrophages sorted by their abundance of bright (acidic) and dim (neutral or non-phagocytic) phagosomes, with gene ranking performed using MAGeCK (Li et al. 2014). In both M0 (resting) and IFNγ-activated states (Fig 1B and 1C, Supplementary Table 1), the screen identified essential phagocytic machinery, including the high-affinity FcγRI (*Fcgr1*), its associated gamma chain (*Fcer1g)*, as well as genes that mediate phagosome acidification including the Ragulator complex (*Lamtor2*, *Lamtor3*, *Raga*, and others) and delivery of the lysosome vacuolar-ATPase by the homotypic fusion and vacuole protein sorting (HOPS) complex (*Vps11*, *Vps16*, *Vps18*). When macrophages were polarized with IFNγ, all members of the NOX2 complex (*Ncf1*, *Cyba*, *Cybb*, *Ncf2*, *Ncf4*), its associated chaperone, Eros (*Bc017643*), and activator PKCδ (*Prckd*), were identified as strong negative regulators of acidic phagosome formation (Fig. 1C and 1D). Surprisingly, the CD47 receptor, SIRPα, was identified as supporting the formation of acidic phagosomes, despite its established role as a “don’t - eat-me” signal (Jaiswal et al. 2009).

**Figure 1.**
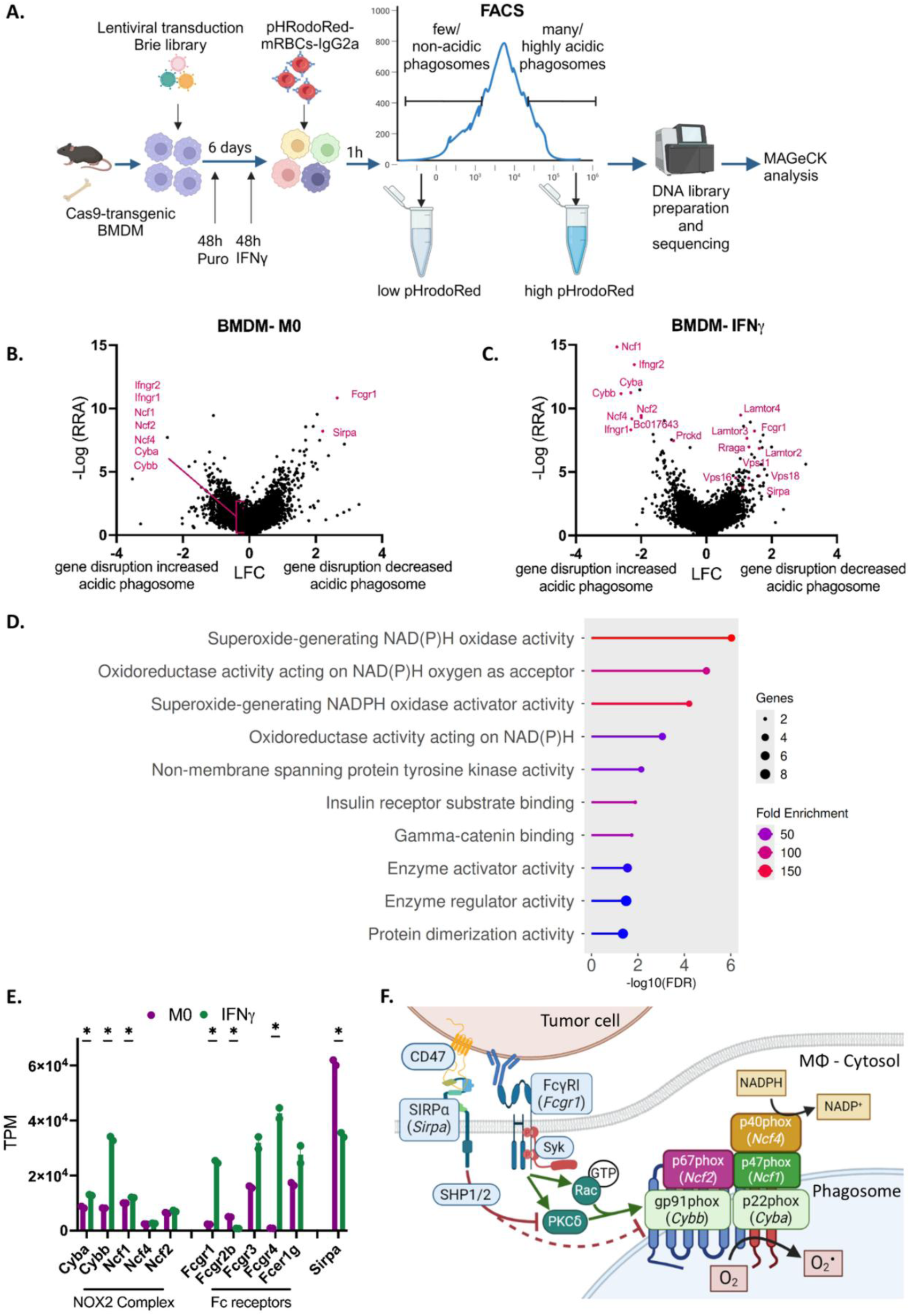
CRISPR-Cas9 whole-genome screen identified the NOX2 complex as an inhibitor of acidic phagosome formation. (A) Workflow for whole-genome CRISPR screen on M0 and IFNγ-polarized macrophages. BMDMs were isolated from Cas9 transgenic mice and transduced with the sgRNA Brie library. Mutant populations were then cultured as M0 or polarized with IFNγ for 48 h, followed by challenge with IgG-coated mRBCs labelled with pHrodo Red-AM. Macrophages with brightly fluorescent acidic phagosomes and those lacking acidic phagosomes were sorted and subjected to sequencing, followed by gene ranking using MAGeCK. Image created in BioRender. (B and C) Volcano plots showing genes responsible for the formation of acidic phagosomes in M0 and IFNγ-polarized BMDMs. The x-axis represents the log fold change (LFC) of gene enrichment, while the y-axis represents the negative logarithm of the Robust Rank Aggregation (log RRA) score, indicating the relative ranking of each gene’s enrichment significance. Known phagocytosis-related genes are highlighted in red. (D) Molecular function terms enriched among the analyzed gene set. Terms are plotted according to statistical significance, expressed as −log10(FDR), with larger values indicating greater significance. Dot size represents the number of genes associated with each term, while dot color indicates fold enrichment. (E) RNA-seq transcript data comparing gene expression of the NOX2 complex members, FcγRs, and *Sirpa* in M0 and IFNγ-polarized BMDMs. Two replicates. Statistical significance was determined by multiple unpaired t-tests. *p < 0.05 (F) Schematic structure model of the NOX2 complex and how CD47-SIRPα and FcγRs regulate its activation. Image created in BioRender.

These findings are consistent with IgG engagement of FcγRs promoting NOX2 assembly and activation on the phagosome, leading to generation of superoxide anion (Balce et al. 2016). NOX2 has been extensively studied in dendritic cells, where superoxide reacts with H^+^ to neutralize phagosome pH and promote lipid peroxidation, leading to the formation of pores in the phagosome membrane to facilitate antigen cross-presentation and antigen preservation (Savina et al. 2006; Rybicka et al. 2010; Dingjan et al. 2016; Embgenbroich and Burgdorf 2018).

Transcriptomic analysis revealed that IFNγ polarization increased the expression of *Cybb*, which encodes gp91-phox, the catalytic subunit of NOX2, along with transcripts encoding activating FcγRs (*Fcgr1*, *Fcgr2*, and *Fcgr3*), while reducing transcripts for the inhibitory FcγR (*Fcgr2b)* and SIRPα (*Sirpa*; Fig. 1E). These findings suggest a model in which IFNγ activation promotes stronger FcγR signaling and NOX2 activation via Syk and PKCδ signaling while lessening CD47-SIRPα-mediated inhibition involving recruitment of the phosphatases SHP-1/2 to the nascent phagosome (Fig. 1F).

### Target cell-associated CD47 suppresses NOX2 activation during ADCP

To determine if target-associated CD47 suppresses NOX2 activation, we measured antibody-dependent ROS using luminol/HRP chemiluminescence during ADCP of target cells with or without CD47 (Fig. 2A). IFNγ polarized wildtype BMDMs exhibited higher luminescence during ADCP of IgG2a opsonized and CD47 blocked A20 cells, whereas *Cybb*^-/-^ BMDMs did not produce ROS (Fig. 2B). Phagocytosis of IgG-opsonized WT or CD47^-/-^ mRBCs revealed that CD47 suppressed antibody-dependent ROS production, with limited effect on ADCP, at a range of 13,750-220,000 IgG per mRBC (Fig. 2C, Supplementary Fig. S1A). Moreover, the amount of ROS generated scaled with the IgG density on the target cell (Fig. 2C). Similar measurements were made for anti-CD20 IgG2a opsonized WT or CD47^-/-^ splenic B cells (SBC) (Fig. 2D, Supplementary Fig. S1B and S1C) and with or without CD47 blockade on A20 lymphoma cells (Fig. 2E). In all cases, target cell CD47 had a modest effect on ADCP (Fig. 2D, 2E) even at sub-pharmacological densities. Thus, for strong surface antigens, IgG2a antibodies readily surpass the CD47 barrier for ADCP; however, CD47 placed a more stringent barrier on antibody-dependent ROS. Moreover, IFNγ polarization potentiated antibody-dependent ROS consistent with the upregulation of *Cybb* (Fig. 1E), but it did not alleviate the CD47 inhibition of antibody-dependent ROS (Fig. 2D and 2E).

**Figure 2.**
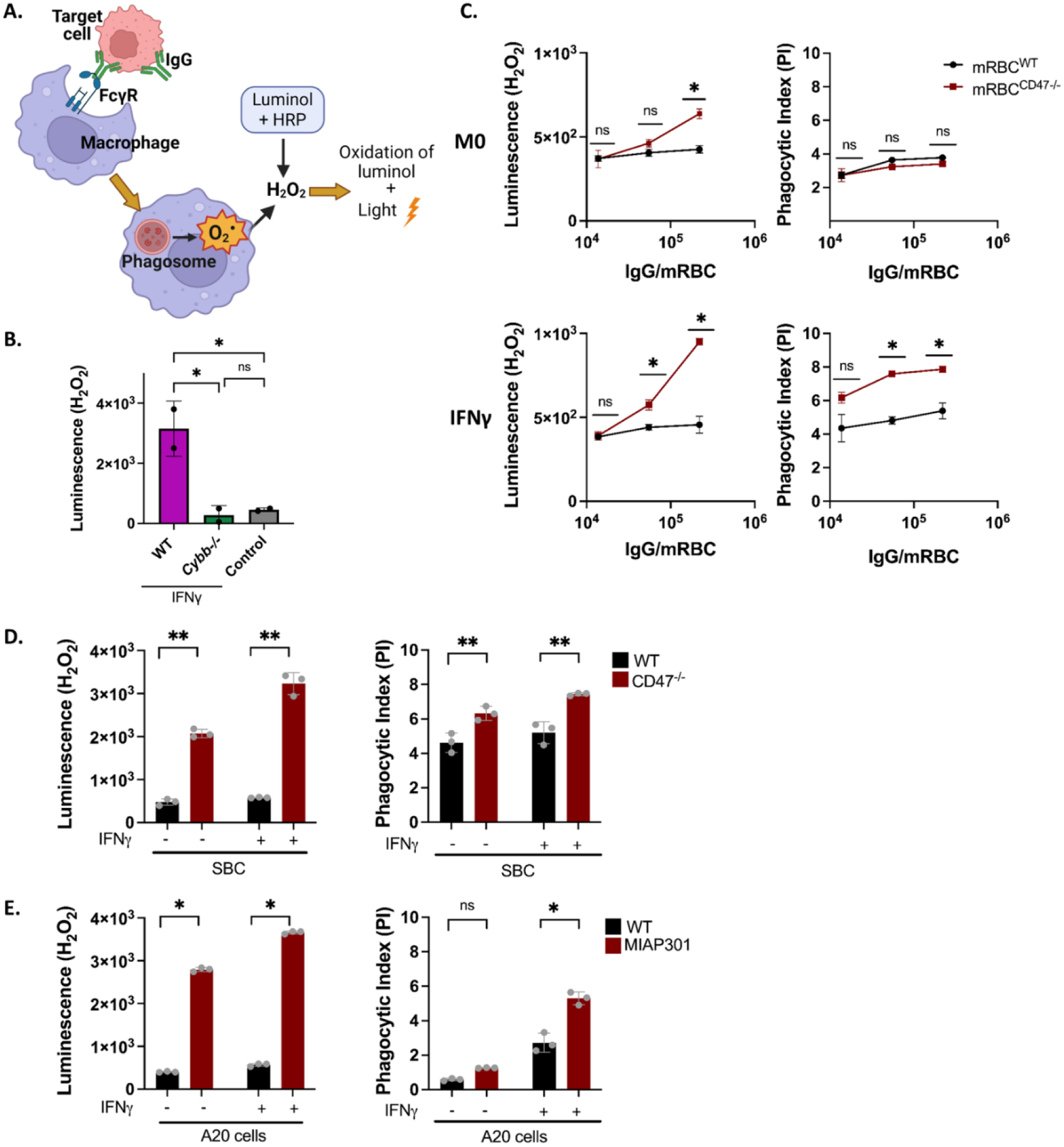
CD47 constrains antibody-dependent ROS in M0 and IFNγ-polarized macrophages. (A) Schematic representation of antibody-dependent ROS production measured by luminol assay. Image created in BioRender. (B) Comparison of antibody-dependent ROS production by IFNγ-polarized WT and *Cybb*^-/-^ BMDMs during ADCP of anti-CD20 IgG2a-opsonized A20 cells and CD47 blocked using MIAP301 (Error bars ± standard deviation; one-way ANOVA followed by Tukey’s multiple comparisons test). (C) Quantification of antibody-dependent ROS production (left panels) and ADCP (right panels) by M0 (top panels) and IFNγ-polarized macrophages (bottom panels) following engulfment of WT (black) or CD47^-/-^ mRBCs (red), measured across increasing concentrations of anti-mRBC-IgG2a (34-3C). Each dot indicates the mean of 3 replicates, representing ∼1,200 BMDM/per replicate. (Error bars ± standard deviation; two-way ANOVA followed by Sidak’s multiple comparisons test within each panel). (D) Antibody-dependent ROS (left panels) and ADCP (right panels) by M0 and IFNγ-polarized macrophages following engulfment of WT (black) or CD47^-/-^ (red) splenic B cells (SBC), measured at a concentration of 1.5 x 10^8^ anti-CD20 IgG2a per SBC. Each dot is an experimental replicate representing ∼1,200 BMDM/per replicate. (Error bars ± standard deviation; two-way ANOVA followed by Sidak’s multiple comparisons test within each panel). (E) Antibody-dependent ROS production (left panels) and ADCP (right panels) by M0 and IFNγ-polarized macrophages following engulfment of WT (black) or MIAP301-treated (red) A20 cells, measured at a concentration of 1.5 x 10^8^ anti-CD20 IgG2a per A20 cell. Each dot is an experimental replicate representing ∼1,200 BMDM/per replicate. (Error bars ± standard deviation; two-way ANOVA followed by Sidak’s multiple comparisons test, as targets and their conditions are independent (D, E)). ns, non-significant; *p < 0.05, **p < 0.01.

### CD47 accelerates phagosome acidification via inhibition of NOX2

To determine if CD47 suppression of NOX2 activity changes phagosome acidification, WT and *Cybb*^-/-^ BMDMs, either M0 or IFNγ-polarized, were incubated with WT or CD47^-/-^ mRBCs opsonized with 10^6^ IgG/mRBC. Ratiometric pH measurements were made by labeling the mRBCs with pHrodo Red, which is brightly fluorescent in the acidic phagolysosome, and AF647-NHS ester, which is pH-insensitive (Fig. 3A). The fluorescence from the mRBCs was then calibrated in buffers at pH 4, 5.5, and 7.4 to enable pH quantification by ratiometric imaging of the pHrodo Red-to-AF647 ratio (Fig. 3B and 3C, Supplementary Fig. S2A). Time-lapse imaging showed that, despite IgG-mRBC with or without CD47 being ingested at similar frequencies, BMDM phagosomes containing mRBC^WT^ acidified rapidly, whereas phagosomes containing mRBC^CD47-/-^ acidified slowly, remaining near neutral pH in IFNγ-polarized macrophages (Fig. 3C, Supplementary Fig. S2B), consistent with their elevated *Cybb,* activating FcγRs and reduced *Sirpa* expression (Fig. 1E). In contrast, *Cybb*^-/-^ BMDMs acidified phagosomes containing WT or CD47^-/-^ mRBC under all conditions (Fig. 3D and 3E), albeit slightly slower than M0-WT BMDMs, demonstrating that CD47 on the target cell suppresses NOX2 to promote rapid phagosome acidification.

**Figure 3.**
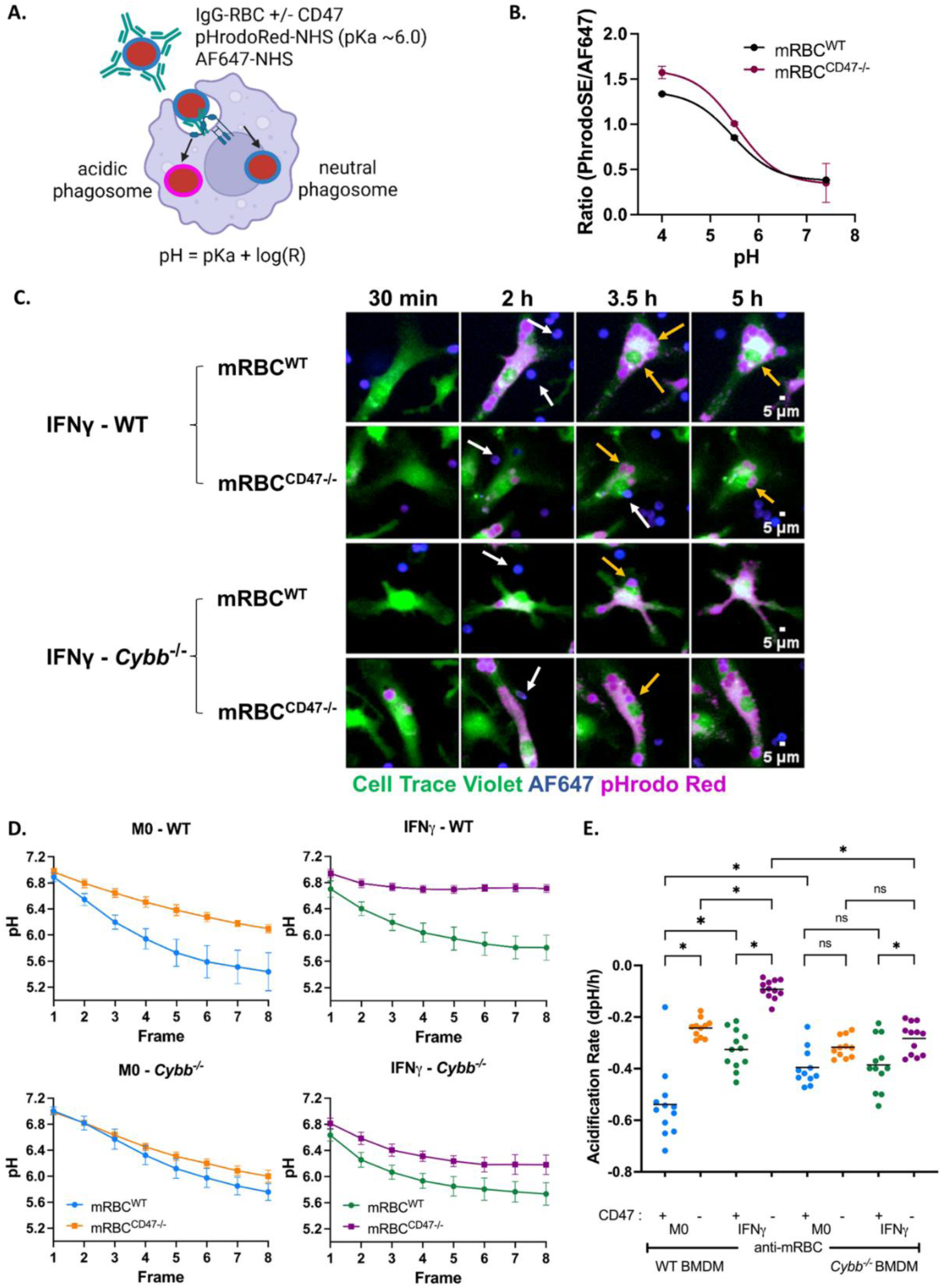
CD47 inhibits NOX2 during ADCP to promote acidification. (A) Schematic of the phagosome pH quantification. IgG-opsonized mRBCs were labeled with a pH-sensitive dye, pHrodo Red, and a pH-insensitive AF647-NHS fluorophore. Image created in BioRender. (B) The pHrodo Red/AF647 fluorescence ratio was used to quantify phagosome acidification and convert fluorescence measurements to pH values. The calibration curve for WT and CD47^-/-^ mRBCs is shown. Each dot represents the ratio of pHrodo Red to AF647 intensities from each of 4 sites/well, representing 1,967-2,257 mRBCs analyzed per condition. Error bars are ± SD of the site means. (C) Time-lapse images of pHrodo Red (magenta) and AF647 (blue) mRBC within phagosomes of WT or *Cybb*^-/-^ BMDM (green) polarized with IFNγ (scale bar, 5µm). White arrows indicate RBCs outside the phagosome, and yellow arrows indicate RBCs inside the phagosome. (D) Phagosome acidification of M0 or IFNγ-polarized macrophages from WT or *Cybb*^-/-^ mice during ADCP of mRBC^WT^ and mRBC^CD47-/-^. Each point is the mean from 4 sites across 3 wells, representing 875-1,400 macrophages analyzed per condition. Error bars are ± SD of the site means. The analysis is representative of 3 independent experiments. (E) Acidification rate calculated from the calibrated pHrodo Red/AF647 ratio inside the phagosome over the first 5 h. Each dot represents the mean pH from three replicate wells (4 sites/well), representing the above-mentioned number of macrophages analyzed per condition. Statistical significance was analyzed by two-way ANOVA with Tukey’s multiple comparisons test. ns, non-significant; *p<0.05, **p<0.01.

### NOX2 activation does not alter ADCP or antibody-mediated killing by macrophages

To test whether NOX2 contributes to ADCP in vivo, we analyzed the depletion of B cells in response to anti-CD20-IgG2a (Fig. 4A). Consistent with our in vitro findings, anti-CD20 B cell depletion in WT and *Cybb*^-/-^ mice had similar responses to anti-CD20 antibodies with or without CD47 blockade. Time-course analysis of peripheral blood revealed similar reductions in B cell frequencies in both genotypes after antibody treatment (Fig. 4C, Supplementary Fig. S3), and this equivalence was maintained in lymphoid compartments, with comparable B cell fractions among CD45^+^CD19^+^ splenocytes (Fig. 4D) and lymph node cells (Fig. 4E). Together, these data indicate that NOX2 is dispensable for ADCP in vivo.

**Figure 4.**
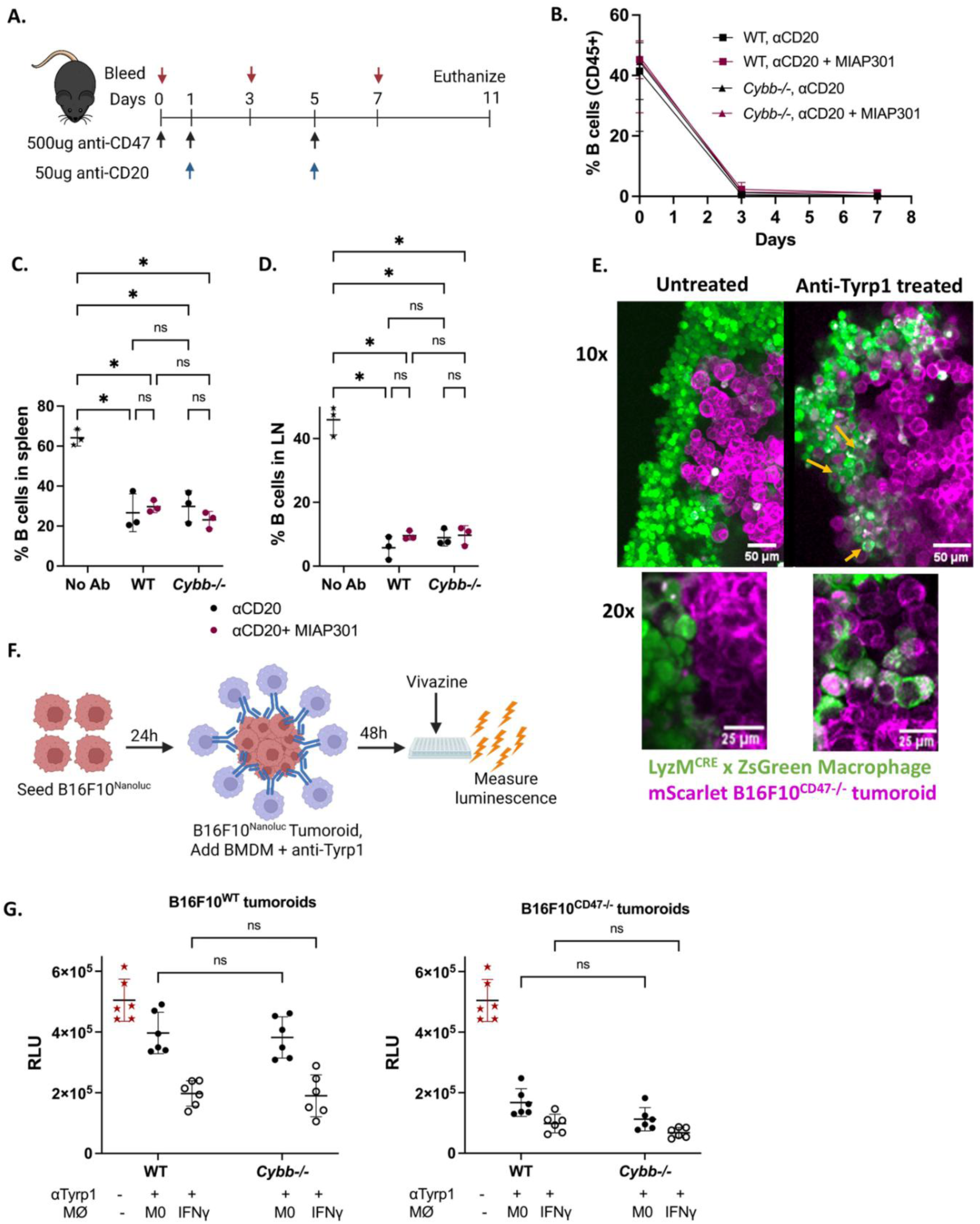
NOX2 is dispensable for macrophage ADCP and does not contribute to antibody-mediated killing of tumoroids. (A) Experimental timeline for B cell depletion in WT and *Cybb*^-/-^ mice. Blood was collected on days 0, 3, and 7. Mice in the treatment groups received anti-CD20 antibody (50 µg per dose on days 1 and 5) either alone or in combination with anti-CD47 antibody (500 µg per dose on days 0, 1, and 5). Mice were euthanized and tissues collected (spleen and inguinal lymph nodes) on day 11. (B) Longitudinal quantification of the percentage of CD19⁺ B cells among CD45⁺ cells in peripheral blood at the indicated time points before and after treatment. Data are shown for n = 3 mice per group. Lines represent group means, and error bars indicate mean ± SD. (C, D) Percentage of remaining B cells out of CD45-positive cells in the spleen (C) and inguinal lymph nodes (LN; D) of WT and *Cybb*^-/-^ mice on day 11 following treatment with anti-CD20 alone or anti-CD20 combined with anti-CD47. Each dot represents an individual mouse. Error bars ± standard deviation; two-way ANOVA with Tukey’s multiple comparison test (E) Confocal images of ADCP of B16F10^CD47-/-^ mScarlet tumoroid by M0 macrophages. LyzM^CRE^ x ZsGreen macrophages are shown in green, and B16F10^CD47-/-^ mScarlet melanoma cells are shown in magenta. 10x scale bar = 50 µm. 20x scale bar = 25 µm. Yellow arrows indicate phagosomes of green macrophages containing magenta B16F10^CD47-/-^ mScarlet cells. (F) Experimental timeline for the in vitro tumoroid depletion assay. B16F10 tumor cells were seeded on day 0, followed by the addition of macrophages, with or without anti-Tyrp1 antibody, on day 1. On day 3, tumoroids were dissociated, vivazine substrate was added, and the luminescence was measured. Image created in BioRender. (G) Quantification of B16F10^WT^ and B16F10^CD47-/-^ tumoroid depletion by WT or *Cybb*^-/-^ BMDMs. Macrophages were either M0 or IFNγ-polarized. Each point represents an individual tumoroid. Error bars indicate the mean ± SD from three replicates across two independent experiments; two-way ANOVA with Tukey’s multiple comparison test. ns, non-significant; *p < 0.05.

We next sought to determine if the increase in ROS generated during anti-Tyrp1-IgG2a-mediated ADCP of CD47-negative tumor cells promoted tumor killing in a high-cellular-density tumoroid context. We used CRISPR to disrupt CD47 in mScarlet-expressing B16F10 melanoma cells, followed by verification by amplicon sequencing of the cut site (Supplementary Fig. S5). As expected from (Dooling et al. 2023), CRISPR-edited mScarlet-B16F10^CD47-/-^ tumoroids cultured with LyzM^CRE^ x ZsGreen BMDM revealed abundant phagosomes within green macrophages during anti-Tyrp1 treatment, but phagosomes were rare in untreated tumoroids (Fig. 4G). To quantify ADCP killing across tumor cell CD47 status, macrophage activation state, and NOX2 function, we developed a 96-well plate luminescence assay. Here, the phagocytic depletion of NanoLuc-expressing B16F10 or B16F10^CD47-/-^ tumoroids provided superior sensitivity to traditional firefly luciferase (Supplementary Fig. S4A and S4B). IFNγ-polarized WT and *Cybb*^-/-^ BMDM partially depleted B16F10^WT^ in the presence of anti-Tyrp1. However, their M0 counterparts had a negligible effect on B16F10^WT^. In contrast, anti-Tyrp1 promoted killing of B16F10^CD47-/-^ by both WT and *Cybb*^-/-^ M0 BMDMs and enabled nearly complete tumor cell elimination by IFNγ-polarized BMDMs of both genotypes Fig. 4G, while the presence of the IFNγ BMDMs has a minimal impact on tumoroid growth in the absence of anti-Tyrp1 (Supplementary Fig. S4C and S4D). In this context, *Cybb*^-/-^ BMDMs depleted B16F10^CD47-/-^ similarly to, or slightly better than, WT BMDMs (Fig. 4G), indicating that NOX2 did not contribute to tumor cell killing and establishing ADCP as the principal mechanism.

### CD47 and NOX2 regulate adaptive immune surveillance and anti-Tyrp1 immune responses

Given the role of NOX2 in antigen presentation in dendritic cells (Savina et al. 2006; Mantegazza et al. 2008) and that removal of CD47 promotes adaptive immune engagement (Dooling et al. 2023), we hypothesized that antigen presentation could be defective in mice lacking NOX2 function during anti-Tyrp1 immunotherapy. To evaluate this, we turned to the B16F10 model, which is an immunogenically ‘cold’ tumor for which therapeutic doses of anti-Tyrp1-IgG2a antibodies are commercially available and is syngeneic in C57BL/6 mice, enabling testing in *Cybb*^-/-^ animals. Here, WT mice bearing B16F10^WT^ or B16F10^CD47⁻/⁻^ tumors were treated with anti-Tyrp1. As expected, anti-Tyrp1 treatment led to ∼36% of the WT mice clearing B16F10^CD47-/-^ but not B16F10^WT^ tumors (Fig. 5A, Supplementary Fig. 6). Of the mice that cleared the initial challenge of B16F10^CD47-/-^ tumors, 3 out of 5 mice rejected a second challenge (Supplementary Fig. S7D). In contrast, *Cybb*^-/-^ mice bearing B16F10^CD47⁻/⁻^ tumors were unable to clear the tumor during anti-Tyrp1 therapy, despite anti-Tyrp1 slowing tumor growth and two animals attaining minimal tumor burden but then relapsing after the conclusion of anti-Tyrp1 therapy (Fig. 5A, Supplementary Fig. S6H). This finding indicated that *Cybb*^-/-^ mice are defective in engaging adaptive immune responses and gaining tumor control in the absence of a therapeutic opsonizing antibody.

**Figure 5.**
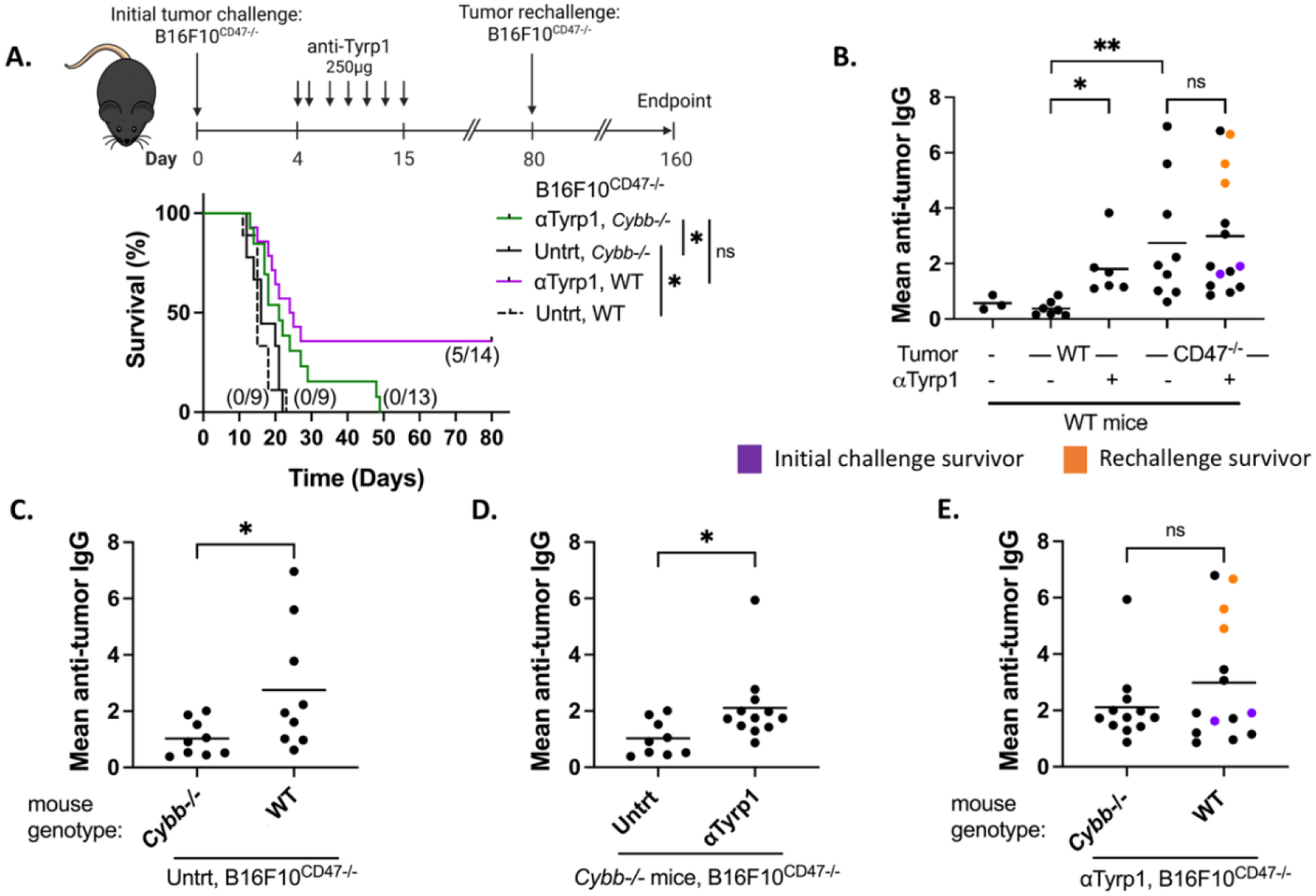
NOX2 is required for anti-Tyrp1-mediated clearance of B16F10^CD47-/-^ tumors, and CD47 prevents immune recognition of B16F10 tumors in a NOX2-dependent pathway. (A) Treatment timeline and survival analysis of WT or *Cybb*^-/-^ mice injected with B16F10^CD47-/-^ melanoma cells. Mice received anti-Tyrp1 antibody (250 μg per dose) on days 4, 5, 7, 9, 11, 13, and 15 after tumor injection. Fractions in parentheses indicate the number of surviving mice over the total number of mice in each cohort. Statistical significance was analyzed by the log-rank (Mantel-Cox) test. (B-E) Quantification of endpoint plasma IgG reactivity against B16F10^CD47-/-^ melanoma cells in the following experimental groups: (B) WT mice, B16F10^WT^ or B16F10^CD47-/-^ tumors, with or without anti-Tyrp1, (C) untreated WT and *Cybb*^-/-^ mice, B16F10^CD47-/-^ tumors, (D) *Cybb*^-/-^ mice, B16F10^CD47-/-^ tumors, with or without anti-Tyrp1 and (E) anti-Tyrp1 treated, WT and *Cybb*^-/-^ mice, B16F10^CD47-/-^ tumors. For all panels, plasma IgG reactivity was assessed by incubating B16F10^CD47-/-^ tumor cells with plasma, followed by detection with fluorescently labeled anti-mouse IgG-AF647. Each dot represents the mean IgG-associated fluorescence intensity for an individual mouse, averaged across three independent plasma-analysis experiments. Purple dots represent mice that survived the initial tumor challenge, whereas orange dots represent mice that survived tumor rechallenge. Statistical significance was analyzed by two-way ANOVA with Tukey’s multiple comparisons test (B) or an unpaired t-test (C, D, and E). ns, non-significant; *p < 0.05; **p < 0.01.

To examine this further, we evaluated the production of anti-B16F10 IgG as a proxy for adaptive immune engagement. Given that the major half-life of anti-Tyrp1 in mouse plasma ranges from 10-15 h (Welt et al. 1987), blood collected 48 h post-injection predominantly reports produced new anti-B16F10 IgG. As expected, B16F10^WT^ tumors did not induce an antibody response, typical of a non-immunogenic tumor. Treatment of B16F10^WT^ tumors with anti-Tyrp1, or removal of CD47 was sufficient to induce anti-tumor IgG production in WT mice (Fig. 5B), but neither correlated with slowed tumor growth (Fig. 5A). Interestingly, anti-Tyrp1 therapy of B16F10^CD47-/-^ did not increase anti-tumor IgG production beyond that seen in untreated B16F10^CD47-/-^ tumors or anti-Tyrp1 treated B16F10^WT^ tumors despite slowing tumor growth and enabling tumor clearance (Fig. 5B). These findings suggest that CD47 removal or anti-Tyrp1 opsonization, allowed immune recognition of the tumor and initiation of an adaptive response, but only the combination of therapeutic opsonizing IgG and the removal of tumor-associated CD47 enables potent tumoricidal activity (Fig. 5B).

In *Cybb*^-/-^ mice, unlike WT mice, CD47 removal was insufficient to produce an anti-tumor IgG response (Fig. 5C), indicating that NOX2 regulation by CD47 is a critical step in immune surveillance. Interestingly, the addition of anti-Tyrp1 produced an antibody response to B16F10^CD47-/-^ tumor cells in *Cybb*^-/-^ mice (Fig. 5D) similar to WT mice (Fig. 5E), despite having a blunted therapeutic effect. Taken together, these data suggest that CD47 constrains NOX2 activation at a basal state, limiting immune surveillance, and that IgG opsonization activates adaptive immune engagement even in the absence of NOX2. From these observations, we hypothesized that strong antigen presentation, supported by NOX2, was needed to generate sufficient T cell activation to produce IFNγ and promote ADCP-mediated killing of the tumor, while the T cell response continues to develop.

### Anti-Tyrp1-mediated clearance of CD47-deficient B16F10 tumors requires IFNγ and CD8 T cells but is inhibited by CD4 T cells

We next sought to determine if T cell activation and IFNγ production were needed to support anti-Tyrp1 killing of B16F10^CD47-/-^ tumors, which have been implicated in effective immune responses driven by tumor opsonizing antibodies (Beck et al. 2024; Wittrup 2017). Indeed, administration of an IFNγ blocking antibody during anti-Tyrp1 therapy abrogated tumor control (Fig. 6A, Supplementary Fig. 7), suggesting that T cell help was needed to support effective ADCP. To determine which T cells were required, we depleted CD4 or CD8 T cells and measured the response of B16F10^CD47-/-^ tumors to anti-Tyrp1 therapy. Strikingly, CD8 T cell depletion abrogated tumor control and anti-tumor IgG production (Fig. 6B and 6C). Conversely, depletion of CD4 T cells provided enhanced tumoricidal responses, attaining a 90% cure rate (Fig. 6B), despite a loss in the production of anti-tumor IgG (Fig. 6C). Surviving CD4 T cell-depleted mice rejected tumor rechallenge in 7 out of 9 mice (Supplementary Fig. S7D), indicating that CD4 T cell removal unmasked strong CD8 T cell contributions to tumor shrinkage and promoted the development of memory against B16F10^CD47-/-^ tumors. Taken together, these data argue that ADCP and robust CD8 T cell activation are essential for anti-Tyrp1 therapy and subsequent induction of protective immunity.

**Figure 6.**
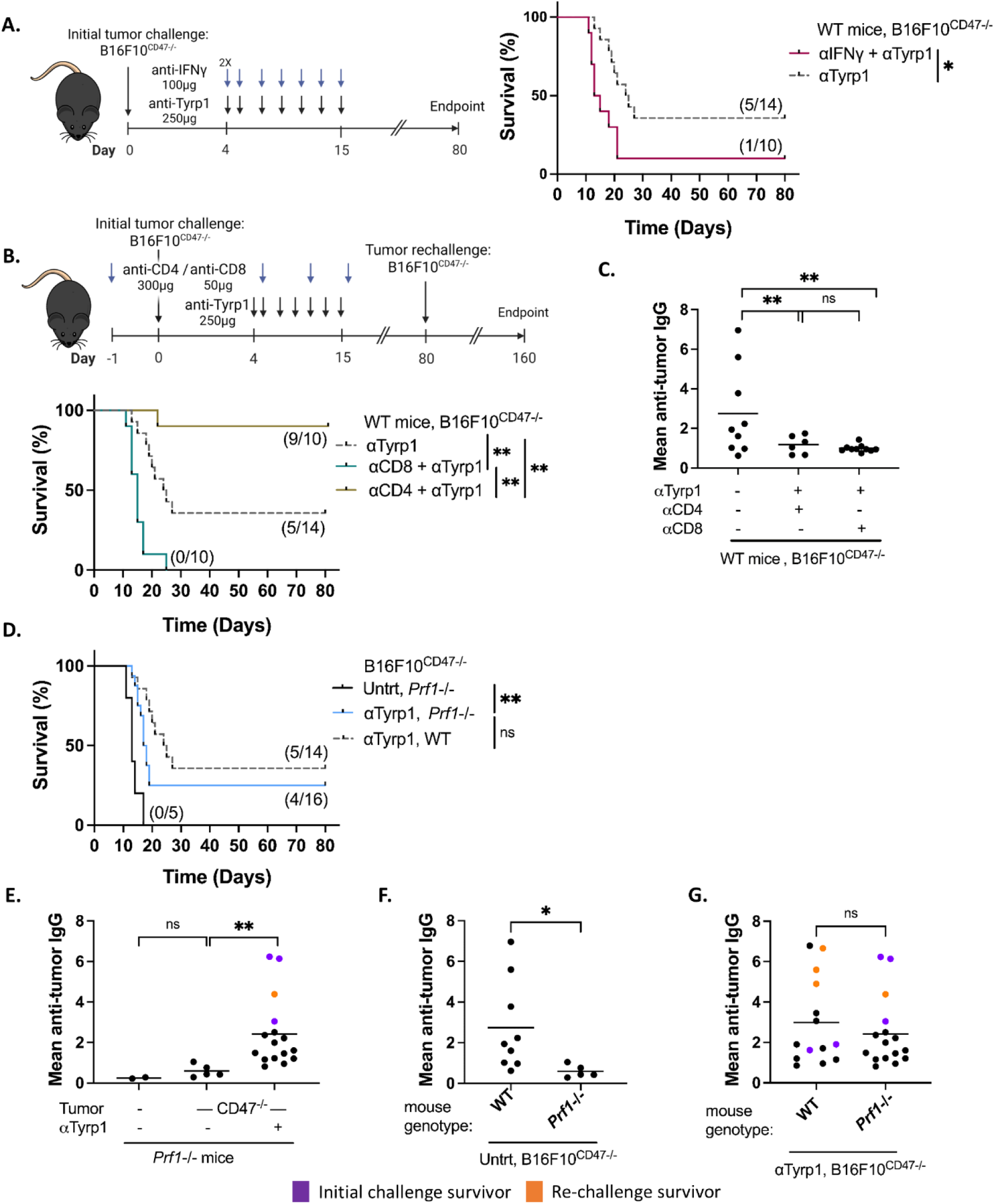
Anti-Tyrp1-mediated tumor clearance requires CD8 T cells and IFNγ but not Perforin and is inhibited by CD4 T cells. (A) Treatment timeline and survival analysis of WT mice injected with B16F10^CD47-/-^ melanoma cells and treated with anti-Tyrp1 antibody with anti-IFNγ blockade. IFNγ was depleted by delivering 200 μg anti-IFNγ antibody on day 4, followed by 100 μg on days 5, 7, 9, 11, 13, and 15, and compared with mice treated with anti-Tyrp1 (grey dashed line). Fractions in parentheses indicate the number of surviving mice over the total number of mice in each cohort. Statistical significance was determined by the log-rank (Mantel-Cox) test. (B) Treatment timeline of WT mice injected with B16F10^CD47-/-^ cells and treated with anti-Tyrp1 antibody along with CD4 or CD8 T cell depletion. CD4 T cells were depleted using 300 μg anti-CD4 antibody and CD8 T cells were depleted using 50 μg anti-CD8 antibody on days -1, 5, 11, and 17, relative to implantation of B16F10^CD47-/-^ cells. Data from the anti-Tyrp1 trial were included (grey dashed line; log-rank (Mantel-Cox) test). (C) Quantification of plasma IgG reactivity against B16F10^CD47-/-^ melanoma cells in CD4- and CD8 T cell-depleted mice. Tumor cells were incubated with plasma and subsequently stained with fluorescently labeled anti-mouse IgG conjugated to AF647. Plasma was taken from blood obtained on day 26 (CD4-depleted mice) or at endpoint (CD8-depleted mice). Two-way ANOVA with Tukey’s multiple comparisons test. (D) Survival analysis of *Prf1*^-/-^ mice injected with B16F10^CD47-/-^ cells and treated with anti-Tyrp1 monotherapy. Data from the anti-Tyrp1 treatment trial were included (grey dashed line; log-rank (Mantel-Cox) test). (E-G) Quantification of endpoint plasma IgG reactivity against B16F10^CD47-/-^ cells in *Prf1*^-/-^ mice. Tumor cells were incubated with plasma and subsequently stained with fluorescently labeled anti-mouse IgG-AF647. Each dot represents the mean IgG-associated fluorescence intensity for an individual mouse, averaged across three independent plasma-analysis experiments. Purple dots represent mice that survived the initial tumor challenge, whereas orange dots represent mice that survived tumor rechallenge. Two-way ANOVA with Tukey’s multiple comparisons test (E); unpaired t-test (F, G). ns, non-significant; *p < 0.05; **p < 0.01.

To determine if cytotoxic T lymphocyte (CTL) function was responsible for anti-Tyrp1 tumor clearance, we examined tumor responses in perforin1-deficient (*Prf1^-/-^)* mice. *Prf1*^-/-^ mice bearing B16F10^CD47-/-^ tumors, treated with anti-Tyrp1 therapy, achieved similar tumor clearance frequencies as WT mice (Fig. 6D), with slightly faster tumor growth rates (Supplementary Fig. S7G and S7H), indicating that the cytolytic function of CTL or NK cells was not required but may contribute to anti-Tyrp1 efficacy against B16F10^CD47-/-^ tumors. Anti-Tyrp1-treated B16F10^CD47-/-^ tumor-bearing *Prf1*^-/-^ mice developed anti-tumor IgG at levels matching those of WT mice (Fig. 6E and 6G). Interestingly, the production of anti-tumor IgG against B16F10^CD47-/-^ tumors without treatment was lost in the *Prf1*^-/-^ mice when compared to WT mice (Fig. 6E and 6F), suggesting that CD47 suppression of immune surveillance may be associated with cytolytic activity. Taken together, these results indicate that macrophage ADCP, driven by opsonizing IgG, and alleviation of CD47 suppression of antigen presentation can drive potent anti-tumor responses.

### MSA-IL-2 or PD-1 blockade improve anti-Tyrp1 tumor clearance in NOX2-deficient, but not WT mice

We reasoned that treatment with IL-2 should support CD8 T cell activation during anti-Tyrp1-therapy by increasing IFNγ production (Beck et al. 2024) and reinforcing an ADCP-CD8 T cell feedback loop. IL-2 is a potent T cell growth factor known to expand tumor-antigen-reactive CD8 T cells and support macrophage-mediated ADCP (Beck et al. 2024). Given the short serum half-life of IL-2 and systemic toxicity at high doses, we used an IL-2 molecule fused to mouse serum albumin (MSA-IL-2) to extend the circulation half-life, enabling persistent low-dose IL-2 (Zhu et al. 2015). MSA-IL-2 alone provided no therapeutic benefit for B16F10^CD47-/-^ tumors (Fig. 7A, Supplementary Fig. S8). Surprisingly, when combined with anti-Tyrp1, MSA-IL-2 did not provide a significant benefit over anti-Tyrp1 alone (Fig. 7A). We then sought to determine if MSA-IL-2 could rescue anti-Tyrp1 therapy in *Cybb*^-/-^ mice by overcoming weak antigen presentation and T cell activation. Here, we found that MSA-IL-2 in combination with anti-Tyrp1 in *Cybb*^-/-^ mice produced excellent B16F10^CD47-/-^ tumor clearance (Fig. 7B), indicating that NOX2 activity can also be inhibitory, likely via release of extracellular ROS suppressing T cell activation and tumor infiltration (Gelderman et al. 2007). To determine if MSA-IL-2 was driving tumor clearance by promoting a CTL response, we next evaluated the combination therapy in *Prf1*^-/-^ mice. Here, MSA-IL-2 + anti-Tyrp1 produced strong tumor clearance (Fig. 7C), confirming that ADCP supported by strong T cell activation and IFNγ production, rather than CTL activity, was likely the principal effector mechanism. Importantly, only one *Prf1*^-/-^ mouse survived re-challenge, indicating the importance of CTL function or durable immunity (Supplementary Fig. S7I and S7J). Together, these data support a model where ADCP, augmented by T cell IFNγ, mediates tumor clearance.

**Figure 7.**
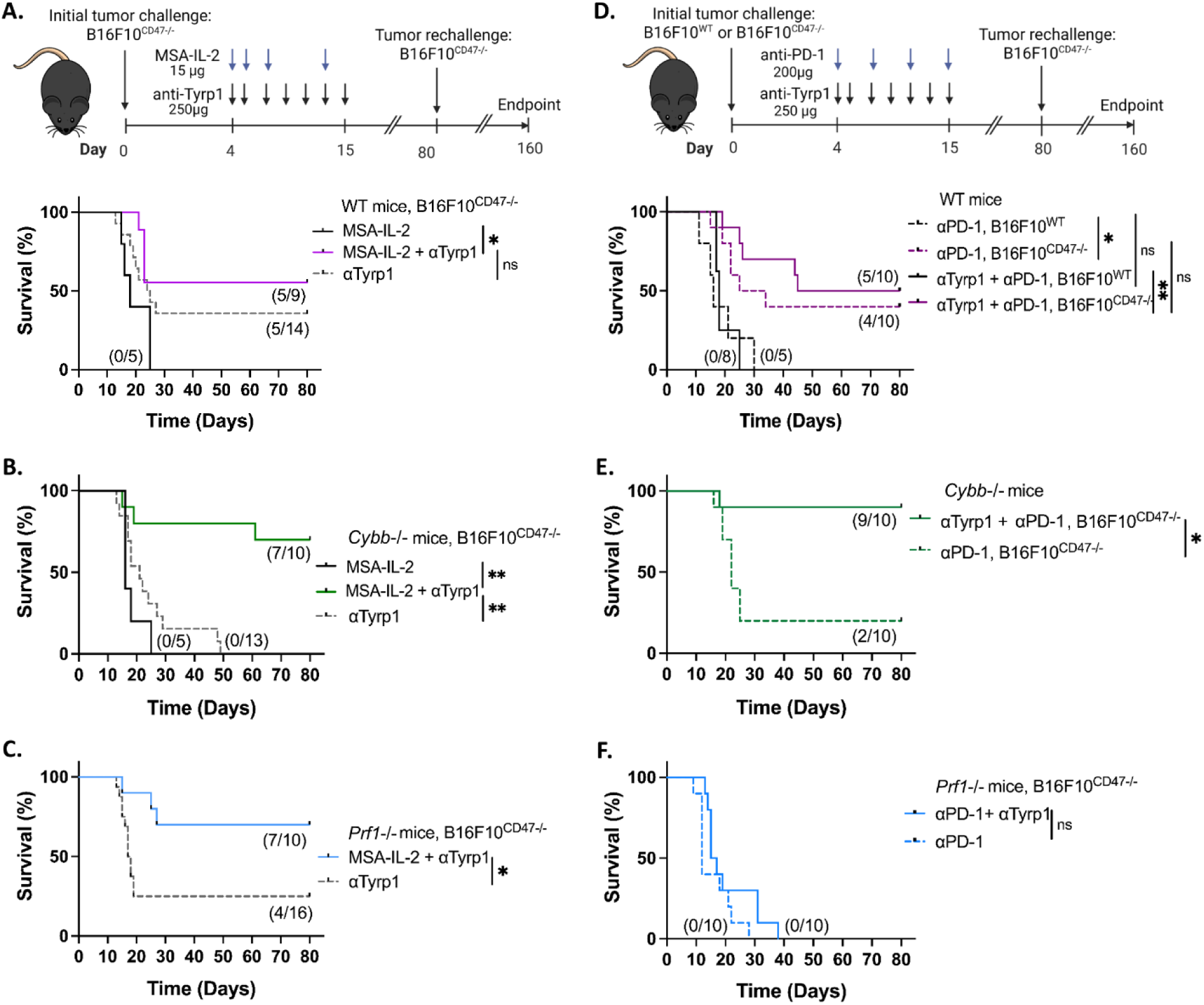
NOX2 restricts and shapes immune responses during MSA-IL-2 and PD-1 blockade enhancement of anti-Tyrp1 therapy of CD47-deficient B16F10 tumors. (A) Treatment timeline and survival plots of WT mice bearing B16F10^CD47-/-^ tumors and treated with MSA-IL-2 or the combination of MSA-IL-2 and anti-Tyrp1 (anti-Tyrp1 alone results shown for comparison, dashed line). The number of surviving mice over the total number of mice in each cohort is shown in parentheses. Statistical significance was determined by the log-rank (Mantel-Cox) test. (B) Survival analysis of *Cybb*^-/-^ mice bearing B16F10^CD47-/-^ tumors treated with MSA-IL-2 or the combination of MSA-IL-2 and anti-Tyrp1, or anti-Tyrp1 alone (dashed line) is included for comparison. Log-rank (Mantel-Cox) test. (C) Survival analysis of *Prf1*^-/-^ mice with B16F10^CD47-/-^ tumors treated with MSA-IL-2 in combination with anti-Tyrp1 antibody or anti-Tyrp1 alone (dashed line) is included for comparison. Log-rank (Mantel-Cox) test. (D) Treatment timeline and survival analysis of WT mice bearing B16F10^WT^ or B16F10^CD47-/-^ tumors and treated with anti-PD-1 monotherapy or the combination of anti-PD-1 and anti-Tyrp1. Solid lines represent combination therapy in mice bearing B16F10^WT^ or B16F10^CD47-/-^ tumors. Log-rank (Mantel-Cox) test. (E) Survival analysis of *Cybb*^-/-^ mice injected with B16F10^CD47-/-^ melanoma cells and treated with anti-PD-1 alone or in combination with anti-Tyrp1 antibody. Log-rank (Mantel-Cox) test. (F) Survival analysis of *Prf1*^-/-^ mice bearing B16F10^CD47-/-^ tumors and treated with anti-PD-1 antibody alone or in combination with anti-Tyrp1 antibody. Log-rank (Mantel-Cox) test. ns, non-significant; *p < 0.05, **p < 0.01.

We reasoned that PD-1 immune checkpoint blockade should also improve anti-Tyrp1 therapy by removing the inhibitory signals during antigen presentation, leading to a more diverse T cell population. We selected a high-affinity Fc null PD-1 blocking antibody, 29F.1A12.1-IgG2c (LALAPG), to more closely approximate pembrolizumab, rather than the more widely used rat isoform or the low-affinity RMP1-14 clone (Bu et al. 2022). In WT mice, B16F10^WT^ tumors were resistant to PD-1 blockade, as found with other, weaker PD-1 blocking antibodies (Curran et al. 2010), and the addition of anti-Tyrp1 provided no additional benefit in tumor control (Fig. 7D, Supplementary Fig. S9A). Unlike during treatment with MSA-IL-2 or with B16F10^WT^ tumors, B16F10^CD47-/-^ tumors were sensitive to PD-1 blockade alone (Fig. 7D, Supplementary Fig. S8B, S9B, and S9D). Although initially unexpected, this result is consistent with our observation that removal of CD47 enabled production of anti-B16F10 antibodies (Fig. 6), highlighting CD47 suppression of surveillance-level T cell activation. Indeed, the combination of anti-Tyrp1 + PD-1 blockade provided minimal improvement in tumor clearance over anti-PD-1 or anti-Tyrp1 alone (Fig. 7D, Supplementary Fig. S9). These results indicate that despite improving the generation of reactive T cells, PD-1 blockade provided only marginal therapeutic improvement, again suggesting an inhibitory pathway.

We then sought to determine if NOX2 regulated the anti-PD-1 + anti-Tyrp1 response. Similar to the combination with MSA-IL-2, anti-Tyrp1 + PD-1 blockade in *Cybb*^-/-^ mice resulted in excellent clearance of B16F10^CD47-/-^ tumors (Fig. 7E, Supplementary Fig. S9E), again pointing to an inhibitory function of NOX2. Moreover, B16F10^CD47-/-^ tumors engrafted in *Cybb*^-/-^ mice responded weakly to PD-1 blockade alone (Fig. 7E), consistent with our observation that in the absence of tumor-associated CD47, NOX2 supported immune surveillance and initial T cell activation. We wondered whetherthe improved anti-tumor response to anti-PD-1 alone or anti-PD-1 + anti-Tyrp1 reflected improved CTL function rather than enhanced ADCP. Indeed, repeating these experiments in *Prf1*^-/-^ mice revealed a complete loss of tumor control during anti-PD-1 alone or anti-PD-1 + anti-Tyrp1 therapies (Fig. 7F). The requirement for perforin1 during PD-1 blockade was consistent with removal of CD47 enabling elevated immune surveillance and primarily CTL. However, the loss of efficacy in the combination treatment suggested a complex interaction between antigen presentation, T cell activation, PD-1 blockade, and ADCP.

### NOX2 is dispensable for durable immunity following MSA-IL-2, but not for PD-1 blockade augmentation of anti-Tyrp1 immunotherapy

Given the striking differences in the *Prf1^-/-^* mice, we sought to gain a better understanding of how MSA-IL-2 and PD-1 blockade supported durable immunity in mice with B16F10^CD47-/-^ tumors. Strikingly, anti-Tyrp1 + MSA-IL-2 therapy survivors from both WT and *Cybb*^-/-^ mice had strong durable immunity (Fig. 8A) that correlated with strong anti-tumor antibody production (Fig. 8C, 8D, and 8E). Notably, this effect required anti-Tyrp1, arguing that ADCP in the presence of MSA-IL-2 drove new antibody production by activating B- and T-responsive mechanisms (Shouse et al. 2024; Ross and Cantrell 2018), raising the possibility that humoral immunity may have contributed to durable immunity in the WT, *Cybb*^-/-^, and *Prf1^-/-^* mice. In contrast, PD-1 blockade conveyed durable immunity as a monotherapy and in the anti-PD-1 + anti-Tyrp1 combination (Fig. 8B), while yielding anti-tumor IgG at levels comparable to anti-Tyrp1-treated wildtype mice (Fig. 8C). Remarkably, despite supporting excellent initial tumor clearance in the *Cybb*^-/-^ mice, the anti-PD-1 + anti-Tyrp1 combination failed to produce durable immunity (Fig. 8B), suggesting a role for NOX2 in the development of high-affinity CTL memory T cells. This observation is consistent with the lack of primary tumor clearance in *Prf1^-/-^* mice (Fig. 7F), supporting the argument that NOX2 plays an important role in the CTL responses. Taken together, these data reveal an underappreciated interplay between tumor-CD47 and the activating and inhibitory effects of NOX2 on T cell activation and ADCP during treatment with opsonizing antibodies.

**Figure 8.**
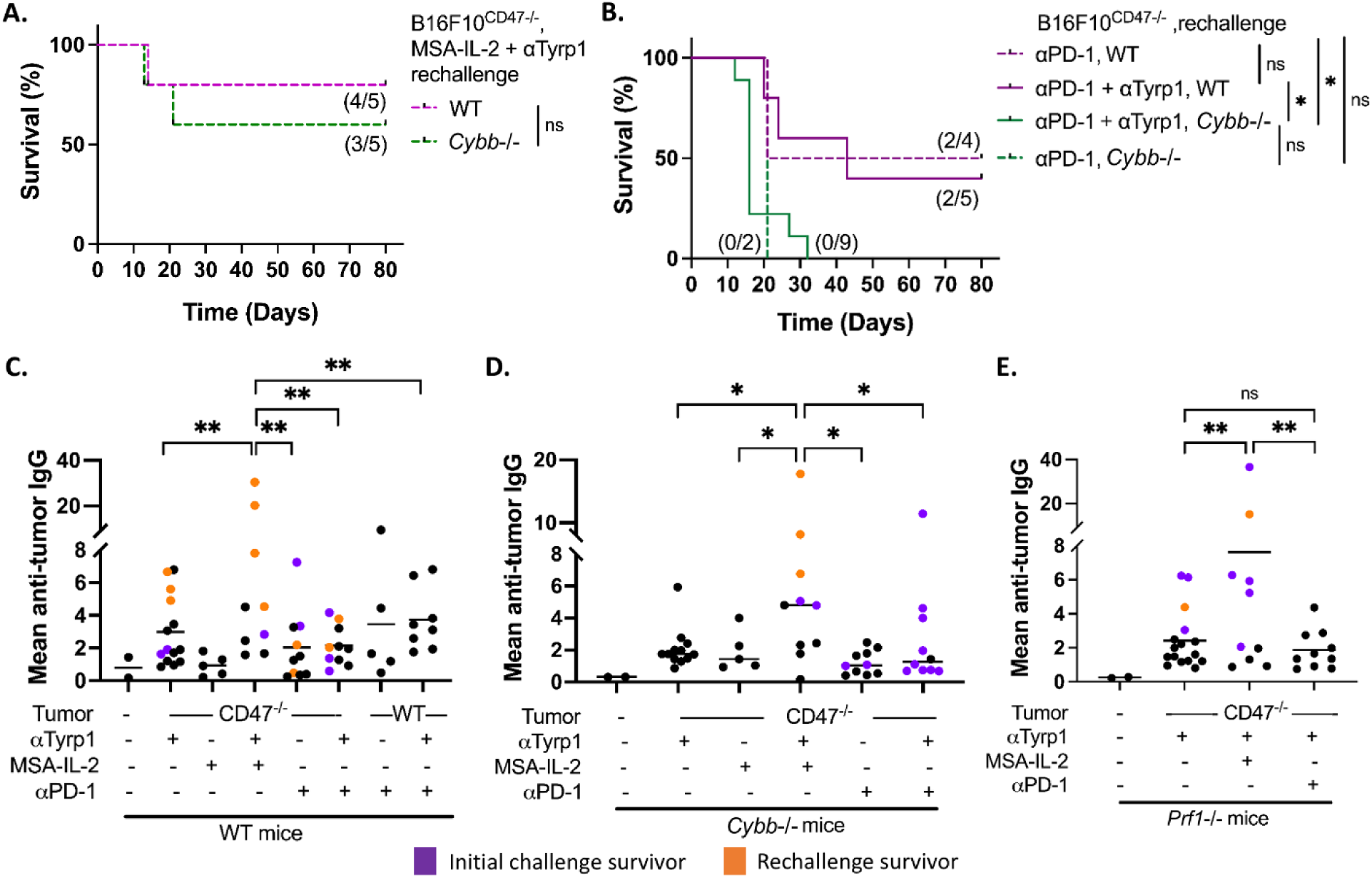
During anti-Tyrp1 therapy of B16F10^CD47-/-^ tumors, MSA-IL-2 supported durable immunity and strong anti-tumor IgG production in WT and *Cybb*^-/-^ mice, whereas PD-1 blockade required NOX2 for durable immunity. (A) Survival analysis of tumor rechallenge of surviving MSA-IL-2 + anti-Tyrp1-treated WT and *Cybb*^-/-^ mice without additional treatment. Log-rank (Mantel-Cox) test. (B) Survival analysis of anti-PD-1 or anti-PD-1 + anti-Tyrp1-treated WT and *Cybb*^-/-^ mice without additional treatment. Log-rank (Mantel-Cox) test. (C-E) Quantification of endpoint plasma IgG reactivity against B16F10^CD47-/-^ cells. Tumor cells were incubated with plasma, followed by staining with fluorescently labeled anti-mouse IgG conjugated to AF647. Each dot represents the mean IgG-associated fluorescence intensity for an individual mouse, averaged across three independent plasma-analysis experiments. Purple dots represent mice that survived the initial tumor challenge, whereas orange dots represent mice that survived tumor rechallenge. Two-way ANOVA with Tukey’s multiple comparisons test. ns, non-significant; *p < 0.05, **p < 0.01.

## Discussion

Our findings reveal an unrecognized function of CD47 in regulating FcγR-driven NOX2 activation during treatment with tumor opsonizing antibody. Specifically, we show that increasing concentrations of opsonizing IgG overcome CD47’s inhibition of phagocytosis, but not CD47’s stringent suppression of NOX2 activation, which ensures rapid phagosome acidification and a weak anti-tumor immune response. Our findings revealed that anti-tumor antibody therapy against CD47-negative tumors required NOX2, CD8 T cells, and IFNγ, arguing that ADCP mediates tumor destruction with support of CD 8 T cells. These results fit a model in which tumor-associated CD47 places a low threshold on phagocytic uptake by inhibiting regulation of the actin cytoskeleton and a high threshold on T cell activation and antigen cross-presentation set by CD47’s regulation of NOX2 activity, with both thresholds lowered by exposure of macrophages to IFNγ. Consistent with this model, augmenting T cell activation with IL-2 or PD-1 blockade restored therapeutic efficacy of anti-Tyrp1 in NOX2-deficient mice and revealed an inhibitory role for NOX2 in the antitumor response.

Together, these findings argue that CD47 serves not only as a “don’t-eat-me” signal, but also as a “don’t-present-me” checkpoint by suppressing IgG-FcγR-NOX2-dependent antigen processing after ADCP. This model aligns with previously published work, demonstrating that opsonizing IgG is essential for tumor-T cell vaccination and adjuvating strategies (Wittrup 2017). Furthermore, it is supported by mounting evidence that targeting CD47-deficient tumors with opsonizing antibodies (Dooling et al. 2023; Beck et al. 2024) or utilizing engineered chimeric antigen receptor-expressing macrophages expressing CD47-blocking nanobodies (Siqi Chen et al. 2025) supports engagement of the adaptive immune system to drive anti-tumor antibody production and CD8 T cell activation.

The requirement for CD8 T cells and IFNγ, but not perforin, for anti-Tyrp1 responses to CD47-negative tumors suggests that CD8 T cells can support antibody-mediated tumor clearance through mechanisms distinct from classic perforin-dependent cytotoxicity. One explanation is that activated CD8 T cells provide IFNγ-mediated help to promote macrophage FcγR-dependent ADCP, establishing a feedback loop between ADCP and T cell activation. This model is consistent with recent evidence that immunotherapy-activated CD8 T cells recruit and promote tumoricidal macrophage states (Van Elsas et al. 2024). Equally important is the finding that depletion of CD4 T cells improved anti-Tyrp1 therapy (Fig. 5). We note that in some instances, mice depleted of CD4 T cells would sometimes have sustained tumor growth followed by rapid shrinkage and disappearance (Fig. 5, Supplementary Fig. S7C), suggesting that CD4 T cells may be beneficial for early tumor control, but may then convert to T regulatory cells that inhibit CD8 T cell and macrophage function.

Future work is needed to identify which specific myeloid subsets govern the FcγR/SIRPα axis to control T cell engagement and effector function. Monocytes and macrophages represent primary candidates, given their high expression of activating and inhibitory FcγRs, SIRPα, and NOX2. In contrast, cDC2s express SIRPα and NOX2, and the inhibitory FcγRIIb, but have limited expression of activating FcγRs. cDC1s, recognized as primary antigen cross-presenting cells, express low levels of activating FcγRs and do not have SIRPα, indicating that additionalmechanisms are likely important for the responses measured here. We verified these expression patterns for B16F10 tumors using publicly available scRNA-seq data (Supplementary Fig. S10; GSM8479167, Yuanhanyu Luo 2024), highlighting that monocytes and macrophages express the preponderance of activating FcγRs, SIRPα, and NOX2 within the tumor. These expression patterns, when compared with our data, are consistent with monocytes and macrophages as essential mediators of ADCP and NOX2-dependent effects, including supportive antigen presentation and inhibitory effects during immune checkpoint blockade. Moreover, these distinct expression patterns suggest that the lack of SIRPα on cDC1 may support immune surveillance of tumor antigens due to a lack of NOX2 inhibition from the CD47-SIRPα pathway and that cDC2s may gain cross-presentation capabilities against CD47-deficient tumors, potentially by lectin- or scavenger receptor-mediated NOX2 activation. Work will be needed to delineate how tumor opsonizing IgG influences which cells contribute to initial and sustained T cell activation.

PD-1 checkpoint blockade or exogenous IL-2 not only rescued anti-Tyrp1-mediated tumor destruction in NOX2-deficient mice but also led to stronger tumor clearance than in WT mice. Our results are consistent with phagosome ROS supporting antigen cross-presentation and initiation of the immune response (Savina et al. 2006; Mantegazza et al. 2008; Rybicka et al. 2010), whereas sustained myeloid-derived oxidants may impose a suppressive threshold on CD8 T cell activation, potentially through peroxynitrite-mediated modification of TCR components (Colligan et al. 2020; X. Chen et al. 2016). This dual role is consistent with the concentration-dependent effects of ROS on T cell signaling, hyperresponsiveness, and apoptosis, and macrophage-derived ROS suppressing IL-2 production, disrupting the reducing environment required for TCR signaling and proliferation, and interfering with cytokine production (Gelderman et al. 2007; Belikov et al. 2015; Bode et al. 2023). Together, these findings support a model in which NOX2-derived ROS promotes antigen processing within the phagosome while imposing a local oxidative brake on lymphocyte function through extracellular or immune-synapse-associated oxidants.

Given the role of NOX2 in regulating phagosome acidification and antigen preservation, CD47-mediated suppression of NOX2 may represent another mechanism by which tumors evade immune recognition. Moreover, our findings identify CD47-dependent inhibition of NOX2 as an important mechanism connecting antibody-mediated phagocytosis to adaptive antitumor immunity, with CD47 influencing not only whether tumor cells are engulfed but also how their antigens are subsequently processed and recognized by the immune system. The requirement for CD8 T cells and IFNγ, but not perforin, further suggests that antibody-mediated tumor clearance depends on cooperative interactions between innate and adaptive immune cells, rather than classical cytotoxicity alone. Finally, the divergent responses to IL-2 and PD-1 blockade reveal that NOX2 may have a complex intersection with antitumor responses and durable immune protection. Defining the responsible phagocyte populations and the mechanisms underlying these opposing effects could inform strategies for combining tumor-opsonizing antibodies with therapies that enhance cancer immunotherapies.

## Acknowledgements

Research reported in this publication was supported by the National Institute of Allergy and Infectious Diseases of the National Institutes of Health under award number U01AI148153 and Cancer Research UK, DRCRPG-May23/100001. The content is solely the responsibility of the authors and does not necessarily represent the official views of the National Institutes of Health. This study was also supported by the Functional Genomic Core Facility (RRID: SCR_023786) and Genomics Sequencing Facility (RRID: SCR_023959) of South Dakota State University.

## Data availability

CRISPR screen files are available in the GEO (GSE353586) database. All data needed to obtain the conclusions in this work are present in the paper or the Supplementary Materials figures or tables. All mouse lines, reagents, and software used are listed in the paper. Any other information will be available upon request.

## References

Balce, Dale R., Joanna M. Rybicka, Catherine J. Greene, Benjamin W. Ewanchuk, and Robin M. Yates. 2016. “Ligation of FCΓR Alters Phagosomal Processing of Protein via Augmentation of NADPH Oxidase Activity.” Traffic 17 (7): 786–802. 10.1111/tra.12396.

Beck, Jan D., Mustafa Diken, Martin Suchan, et al. 2024. “Long-Lasting mRNA-Encoded Interleukin-2 Restores CD8+ T Cell Neoantigen Immunity in MHC Class I-Deficient Cancers.” Cancer Cell 42 (4): 568–582.e11. 10.1016/j.ccell.2024.02.013.

Belikov, Aleksey V., Burkhart Schraven, and Luca Simeoni. 2015. “T Cells and Reactive Oxygen Species.” Journal of Biomedical Science 22 (1): 85. 10.1186/s12929-015-0194-3.

Bess, Shelby N., Matthew J. Igoe, and Timothy J. Muldoon. 2025. “The Physiological and Therapeutic Role of CD47 in Macrophage Function and Cancer.” Immunological Investigations 54 (1): 112–46. 10.1080/08820139.2024.2415409.

Bode, Kevin, Mathias Hauri-Hohl, Vincent Jaquet, and Heiko Weyd. 2023. “Unlocking the Power of NOX2: A Comprehensive Review on Its Role in Immune Regulation.” Redox Biology 64 (August): 102795. 10.1016/j.redox.2023.102795.

Bray, Mark-Anthony, Martha S. Vokes, and Anne E. Carpenter. 2015. “Using CellProfiler for Automatic Identification and Measurement of Biological Objects in Images.” Current Protocols in Molecular Biology 109 (1). 10.1002/0471142727.mb1417s109.

Bu, Melissa T., Long Yuan, Alyssa N. Klee, and Gordon J. Freeman. 2022. “A Comparison of Murine PD-1 and PD-L1 Monoclonal Antibodies.” Monoclonal Antibodies in Immunodiagnosis & Immunotherapy 41 (4): 202–9. 10.1089/mab.2021.0068.

Canton, Johnathan, Hanna Blees, Conor M. Henry, et al. 2021. “The Receptor DNGR-1 Signals for Phagosomal Rupture to Promote Cross-Presentation of Dead-Cell-Associated Antigens.” Nature Immunology 22 (2): 140–53. 10.1038/s41590-020-00824-x.

Cao, Xu, Jing Chen, Bolei Li, et al. 2022. “Promoting Antibody-Dependent Cellular Phagocytosis for Effective Macrophage-Based Cancer Immunotherapy.” Science Advances 8 (11): eabl9171. 10.1126/sciadv.abl9171.

Chen, Shanze, Abdullah F. U. H. Saeed, Quan Liu, et al. 2023. “Macrophages in Immunoregulation and Therapeutics.” Signal Transduction and Targeted Therapy 8 (1): 207. 10.1038/s41392-023-01452-1.

Chen, Siqi, Yingyu Wang, Jessica Dang, et al. 2025. “CAR Macrophages with Built-In CD47 Blocker Combat Tumor Antigen Heterogeneity and Activate T Cells via Cross-Presentation.” Nature Communications 16 (1): 4069. 10.1038/s41467-025-59326-9.

Chen, Xinfeng, Mengjia Song, Bin Zhang, and Yi Zhang. 2016. “Reactive Oxygen Species Regulate T Cell Immune Response in the Tumor Microenvironment.” Oxidative Medicine and Cellular Longevity 2016 (1): 1580967. 10.1155/2016/1580967.

Clement, Kendell, Holly Rees, Matthew C. Canver, et al. 2019. “CRISPResso2 Provides Accurate and Rapid Genome Editing Sequence Analysis.” Nature Biotechnology 37 (3): 224–26. 10.1038/s41587-019-0032-3.

Colligan, Sean H., Stephanie L. Tzetzo, and Scott I. Abrams. 2020. “Myeloid-Driven Mechanisms as Barriers to Antitumor CD8+ T Cell Activity.” Molecular Immunology 118 (February): 165–73. 10.1016/j.molimm.2019.12.012.

Curran, Michael A., Welby Montalvo, Hideo Yagita, and James P. Allison. 2010. “PD-1 and CTLA-4 Combination Blockade Expands Infiltrating T Cells and Reduces Regulatory T and Myeloid Cells within B16 Melanoma Tumors.” Proceedings of the National Academy of Sciences 107 (9): 4275–80. 10.1073/pnas.0915174107.

Dingjan, Ilse, Daniëlle Rj Verboogen, Laurent M. Paardekooper, et al. 2016. “Lipid Peroxidation Causes Endosomal Antigen Release for Cross-Presentation.” Scientific Reports 6 (1): 22064. 10.1038/srep22064.

Doench, John G., Ella Hartenian, Daniel B. Graham, et al. 2014. “Rational Design of Highly Active sgRNAs for CRISPR-Cas9-Mediated Gene Inactivation.” Nature Biotechnology 32 (12): 1262–67. 10.1038/nbt.3026.

Dooling, Lawrence J., Jason C. Andrechak, Brandon H. Hayes, et al. 2023. “Cooperative Phagocytosis of Solid Tumours by Macrophages Triggers Durable Anti-Tumour Responses.” Nature Biomedical Engineering 7 (9): 1081–96. 10.1038/s41551-023-01031-3.

Embgenbroich, Maria, and Sven Burgdorf. 2018. “Current Concepts of Antigen Cross-Presentation.” Frontiers in Immunology 9 (July): 1643. 10.3389/fimmu.2018.01643.

Gelderman, Kyra A., Malin Hultqvist, Angela Pizzolla, et al. 2007. “Macrophages Suppress T Cell Responses and Arthritis Development in Mice by Producing Reactive Oxygen Species.” Journalof Clinical Investigation 117 (10): 3020–28. 10.1172/JCI31935.

Hirayama, Daisuke, Tomoya Iida, and Hiroshi Nakase. 2017. “The Phagocytic Function of Macrophage-Enforcing Innate Immunity and Tissue Homeostasis.” International Journal of Molecular Sciences 19 (1): 92. 10.3390/ijms19010092.

Hodny, Zdenek, Milan Reinis, Sona Hubackova, Pavla Vasicova, and Jiri Bartek. 2016. “Interferon Gamma/NADPH Oxidase Defense System in Immunity and Cancer.” OncoImmunology 5 (2): e1080416. 10.1080/2162402X.2015.1080416.

Hossain, Md. Saddam, Jason G. Kerkvliet, and Adam D. Hoppe. 2023. “Whole Genome CRISPR Screening Strategy to Identify Genes Contributing to SARS-CoV-2 Spike and VSV-G Mediated Entry.” Journal of Medical Virology 95 (9): e29087. 10.1002/jmv.29087.

Hussain, Khiyam, Mark S. Cragg, and Stephen A. Beers. 2021. “Remodeling the Tumor Myeloid Landscape to Enhance Antitumor Antibody Immunotherapies.” Cancers 13 (19): 4904. 10.3390/cancers13194904.

Jaiswal, Siddhartha, Catriona H. M. Jamieson, Wendy W. Pang, et al. 2009. “CD47 Is Upregulated on Circulating Hematopoietic Stem Cells and Leukemia Cells to Avoid Phagocytosis.” Cell 138 (2): 271–85. 10.1016/j.cell.2009.05.046.

Kim, D., J. Wang, S. B. Willingham, R. Martin, G. Wernig, and I. L. Weissman. 2012. “Anti-CD47 Antibodies Promote Phagocytosis and Inhibit the Growth of Human Myeloma Cells.” Leukemia 26 (12): 2538–45. 10.1038/leu.2012.141.

Li, Wei, Han Xu, Tengfei Xiao, et al. 2014. “MAGeCK Enables Robust Identification of Essential Genes from Genome-Scale CRISPR/Cas9 Knockout Screens.” Genome Biology 15 (12): 554. 10.1186/s13059-014-0554-4.

Majeti, Ravindra, Mark P. Chao, Ash A. Alizadeh, et al. 2009. “CD47 Is an Adverse Prognostic Factor and Therapeutic Antibody Target on Human Acute Myeloid Leukemia Stem Cells.” Cell 138 (2): 286–99. 10.1016/j.cell.2009.05.045.

Mantegazza, Adriana R., Ariel Savina, Mónica Vermeulen, et al. 2008. “NADPH Oxidase Controls Phagosomal pH and Antigen Cross-Presentation in Human Dendritic Cells.” Blood 112 (12): 4712–22. 10.1182/blood-2008-01-134791.

Marshall, Michael J. E., Richard J. Stopforth, and Mark S. Cragg. 2017. “Therapeutic Antibodies: What Have We Learnt from Targeting CD20 and Where Are We Going?” Frontiers in Immunology 8 (October): 1245. 10.3389/fimmu.2017.01245.

Mirza, Wajahat, Sundas Dadan, Eshan Ahmad, Ayesha Junaid, and Tariq Mahmood Satti. 2025. “Clinical Outcomes and Safety of CD47-Targeted Immunotherapies across Hematologic Malignancies: A Systematic Review of Monoclonal Antibodies and Fusion Proteins in Combination Strategies.” Clinical and Experimental Medicine 25 (1): 360. 10.1007/s10238-025-01915-5.

Osorio, Juan C., Patrick Smith, David A. Knorr, and Jeffrey V. Ravetch. 2023. “The Antitumor Activities of Anti-CD47 Antibodies Require Fc-FcγR Interactions.” Cancer Cell 41 (12): 2051–2065.e6. 10.1016/j.ccell.2023.10.007.

Ross, Sarah H., and Doreen A. Cantrell. 2018. “Signaling and Function of Interleukin-2 in T Lymphocytes.” Annual Review of Immunology 36 (1): 411–33. 10.1146/annurev-immunol-042617-053352.

Rybicka, Joanna M., Dale R. Balce, Morgan F. Khan, Regina M. Krohn, and Robin M. Yates. 2010. “NADPH Oxidase Activity Controls Phagosomal Proteolysis in Macrophages through Modulation of the Lumenal Redox Environment of Phagosomes.” Proceedings of the National Academy of Sciences 107 (23): 10496–501. 10.1073/pnas.0914867107.

Savina, Ariel, Carolina Jancic, Stephanie Hugues, et al. 2006. “NOX2 Controls Phagosomal pH to Regulate Antigen Processing during Crosspresentation by Dendritic Cells.” Cell 126 (1): 205–18. 10.1016/j.cell.2006.05.035.

Shouse, Acacia N., Kathryn M. LaPorte, and Thomas R. Malek. 2024. “Interleukin-2 Signaling in the Regulation of T Cell Biology in Autoimmunity and Cancer.” Immunity 57 (3): 414–28. 10.1016/j.immuni.2024.02.001.

Stewart, Sheila A., Derek M. Dykxhoorn, Deborah Palliser, et al. 2003. “Lentivirus-Delivered Stable Gene Silencing by RNAi in Primary Cells.” *RNA (New York*, N.Y.) 9 (4): 493–501. 10.1261/rna.2192803.

Swanson, Joel A., and Adam D. Hoppe. 2004. “The Coordination of Signaling during Fc Receptor-Mediated Phagocytosis.” Journal of Leukocyte Biology 76 (6): 1093–103. 10.1189/jlb.0804439.

Trepte, Philipp, Sabrina Kruse, Simona Kostova, et al. 2018. “LuTHy: A Double-Readout Bioluminescence-Based Two-Hybrid Technology for Quantitative Mapping of Protein-Protein Interactions in Mammalian Cells.” Molecular Systems Biology 14 (7): e8071. 10.15252/msb.20178071.

Tsao, Li-Chung, Erika J. Crosby, Timothy N. Trotter, et al. 2019. “CD47 Blockade Augmentation of Trastuzumab Antitumor Efficacy Dependent on Antibody-Dependent Cellular Phagocytosis.” JCI Insight 4 (24): e131882. 10.1172/jci.insight.131882.

Van Elsas, Marit J., Jim Middelburg, Camilla Labrie, et al. 2024. “Immunotherapy-Activated T Cells Recruit and Skew Late-Stage Activated M1-like Macrophages That Are Critical for Therapeutic Efficacy.” Cancer Cell 42 (6): 1032–1050.e10. 10.1016/j.ccell.2024.04.011.

Welt, S., M. J. Mattes, R. Grando, et al. 1987. “Monoclonal Antibody to an Intracellular Antigen Images Human Melanoma Transplants in Nu/Nu Mice.” Proceedings of the National Academy of Sciences of the United States of America 84 (12): 4200–4204. JSTOR.

Willingham, Stephen B., Jens-Peter Volkmer, Andrew J. Gentles, et al. 2012. “The CD47-Signal Regulatory Protein Alpha (SIRPa) Interaction Is a Therapeutic Target for Human Solid Tumors.” Proceedings of the National Academy of Sciences of the United States of America 109 (17): 6662–67. 10.1073/pnas.1121623109.

Wittrup, K. Dane. 2017. “Antitumor Antibodies Can Drive Therapeutic T Cell Responses.” Trends in Cancer 3 (9): 615–20. 10.1016/j.trecan.2017.07.001.

Wöhner, Miriam, and Falk Nimmerjahn. 2025. “Cytotoxic IgG: Mechanisms, Functions, and Applications.” Immunity 58 (6): 1378–95. 10.1016/j.immuni.2025.05.008.

Zhu, Eric F., Shuning A. Gai, Cary F. Opel, et al. 2015. “Synergistic Innate and Adaptive Immune Response to Combination Immunotherapy with Anti-Tumor Antigen Antibodies and Extended Serum Half-Life IL-2.” Cancer Cell 27 (4): 489–501. 10.1016/j.ccell.2015.03.004.

