## Supplemental Data for "Antibody-Fcγ receptor-driven NOX2 activation during phagocytosis of tumor cells is suppressed by CD47, preventing CD8 T cell engagement and immunotherapeutic responses"

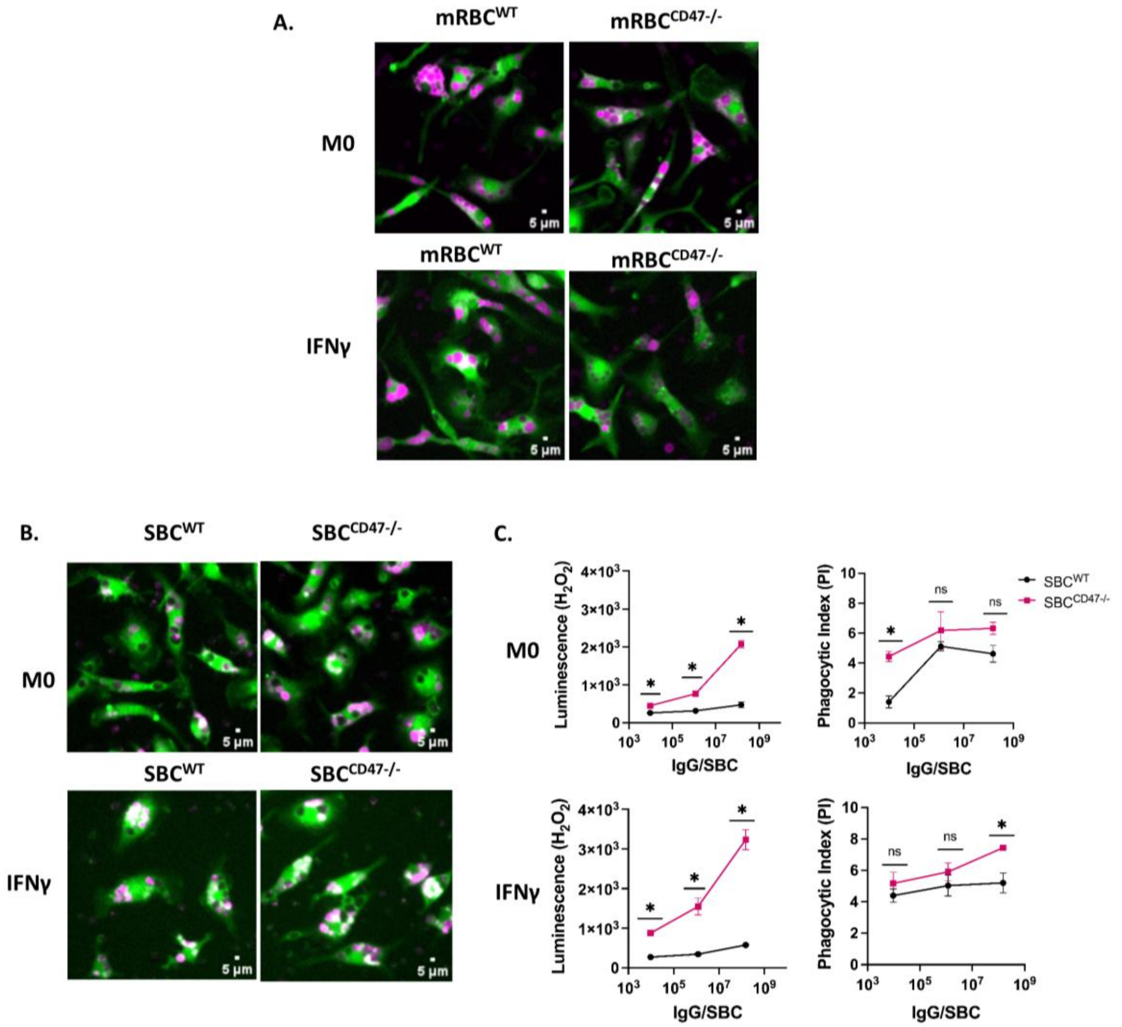


**Figure S1. Comparison of ADCP and ADROS between WT and CD47$^{-/-}$ target cells.**

(A) High-content fluorescence microscopy images of M0 or IFNγ-polarized macrophages following ADCP of WT or CD47$^{-/-}$ mouse red blood cells (mRBC), corresponding to the quantification shown in Fig. 2B. Macrophages (green) and mouse red blood cells (magenta; labeled with pHrodo Red-SE). Images are representative of 3 replicates. Scale bar: 5µm.

(B) High-content fluorescence microscopy images of M0 or IFNγ-polarized macrophages following ADCP of WT or CD47$^{-/-}$ splenic B cells (SBC). Macrophages (green) and splenic B cells (magenta; labeled with pHrodo Red-SE). Images are representative of 3 replicates. Scale bar: 5µm.

(C) Quantification of antibody-dependent ROS production (left panels) and ADCP (right panels) by M0 (top panels) and IFNγ-polarized (bottom panels) macrophages following engulfment of WT (black) or CD47$^{-/-}$ (red) SBCs, measured across increasing concentrations of anti-CD20-IgG2a. Each dot indicates the mean of 3 replicates, representing ~1,200 BMDM/per replicate. (Error bars ± standard deviation; two-way ANOVA followed by Sidak's multiple comparisons test within each panel).

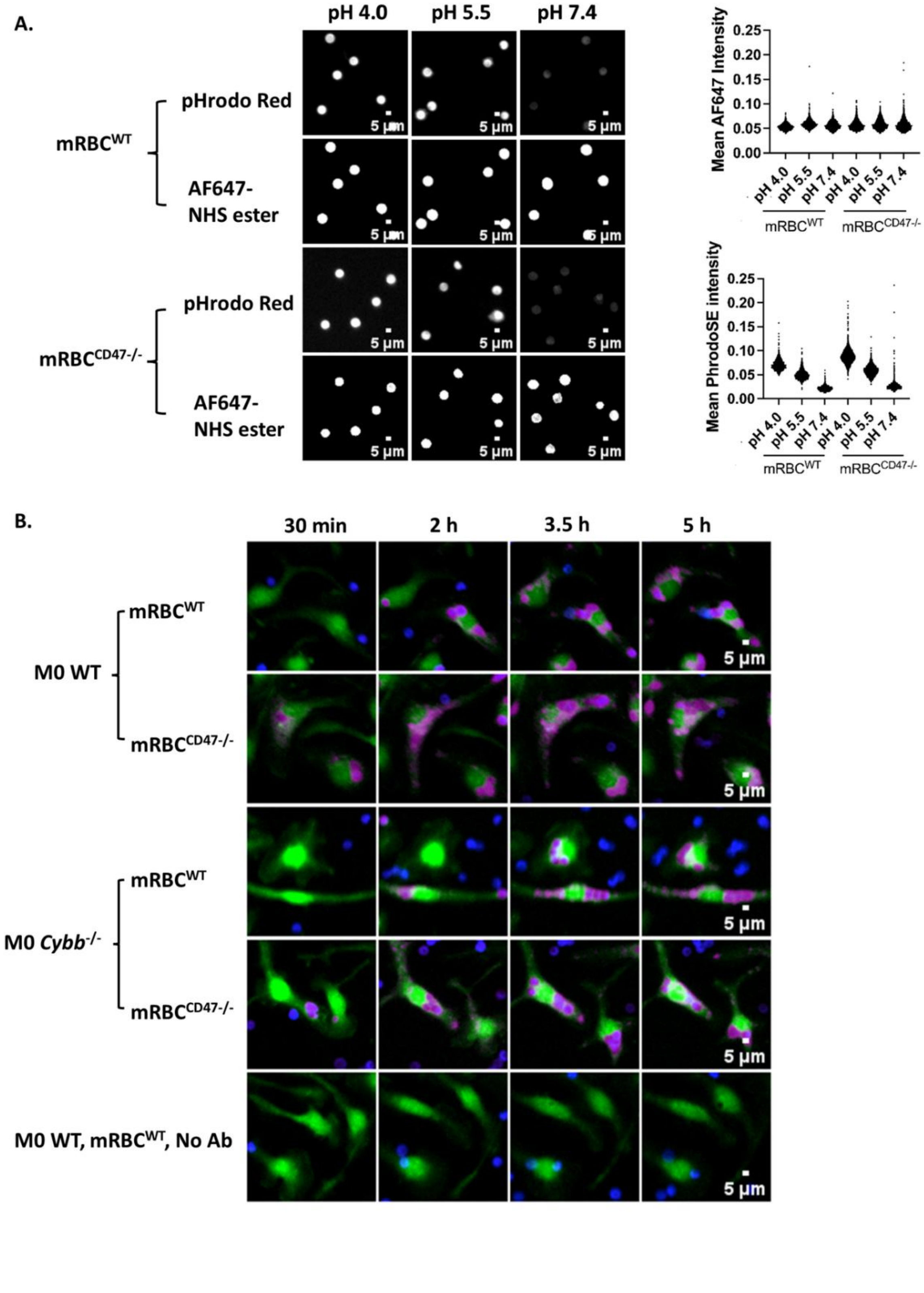
A.
pH 4.0
pH 5.5
pH 7.4
mRBC^WT
pHrodo Red
AF647-
NHS ester
mRBC^CD47-/-
pHrodo Red
AF647-
NHS ester
5 μm
Mean AF647 Intensity
0.25
0.20
0.15
0.10
0.05
0.00
pH 4.0
pH 5.5
pH 7.4
mRBC^WT
mRBC^CD47-/-
Mean PhrodoSE intensity
B.
30 min
2 h
3.5 h
5 h
M0 WT
mRBC^WT
mRBC^CD47-/-
M0 Cybb^-/-
M0 WT, mRBC^WT, No Ab
5 μm

**Figure S2. Calibration of pHrodo Red- and AF647-labeled RBCs in buffers at different pH and visualization of macrophage phagosome acidification in resting-stage macrophages.**

(A) Calibration of labeled mRBCs for phagosomal pH measurement. Representative high-content fluorescence images from the calibration of the WT or $CD47^{-/-}$ mRBCs labeled with the pH-sensitive dye, pHrodo Red, and the pH-insensitive dye, AF647-NHS ester, and incubated in calibration buffers at pH 4.0, 5.5, and 7.4. Scale bar: 5µm. Mean AF-647 or pHrodo fluorescence intensities were quantified across 4 sites/well, representing 1,967-2,257 mRBCs analyzed per condition.

(B) Time-lapse image montages showing phagocytosis and phagosomal acidification over 5 h. WT or $CD47^{-/-}$ mRBCs were labeled with pHrodo Red-SE (magenta) and AF647-NHS ester (blue) before opsonization with mIgG2a (clone 343-C). Resting WT or *Cybb*$^{-/-}$ macrophages were stained with Cell Trace Violet (green). Scale bar: 5µm.

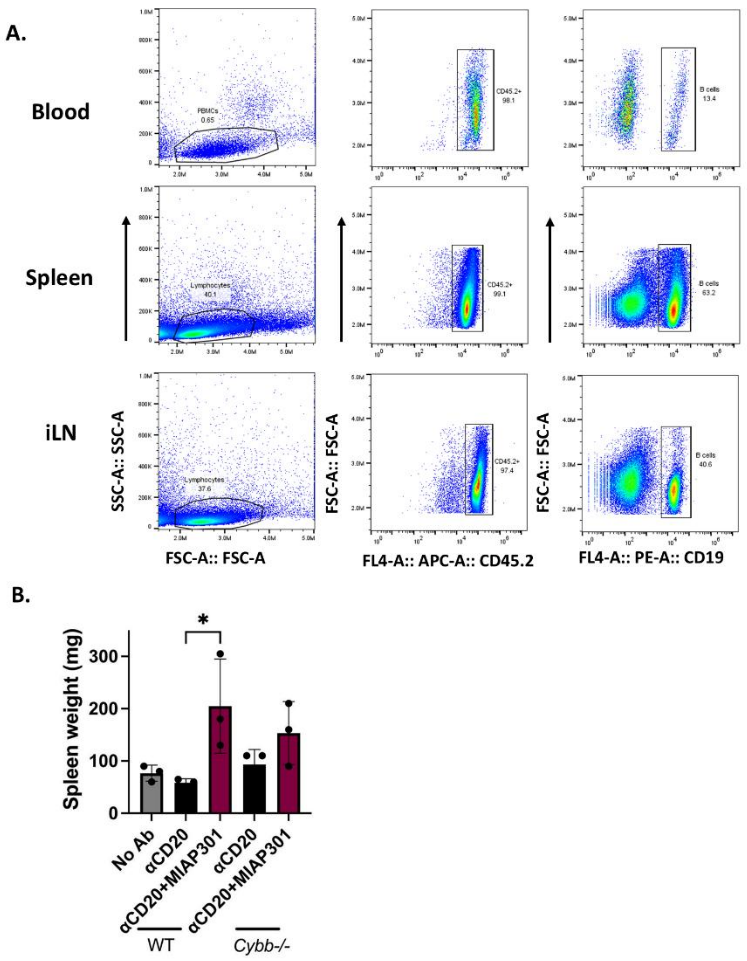


**Figure S3. Gating strategy for CD45.2$^{+}$ CD19$^{+}$ B cells across tissues and effect of CD47 blockade on spleen weight in WT and Cybb$^{-/-}$ mice.**

(A) Flow cytometry gating strategy used for the quantification of CD19-positive B cells among CD45-positive cells remaining in the blood, spleen, or inguinal lymph node following depletion.

(B) Spleen weights (mg) of WT or *Cybb*$^{-/-}$ mice across the indicated treatment groups. (n = 3 per group, error bars ± standard deviation, ordinary one-way ANOVA with Tukey's multiple comparison test)

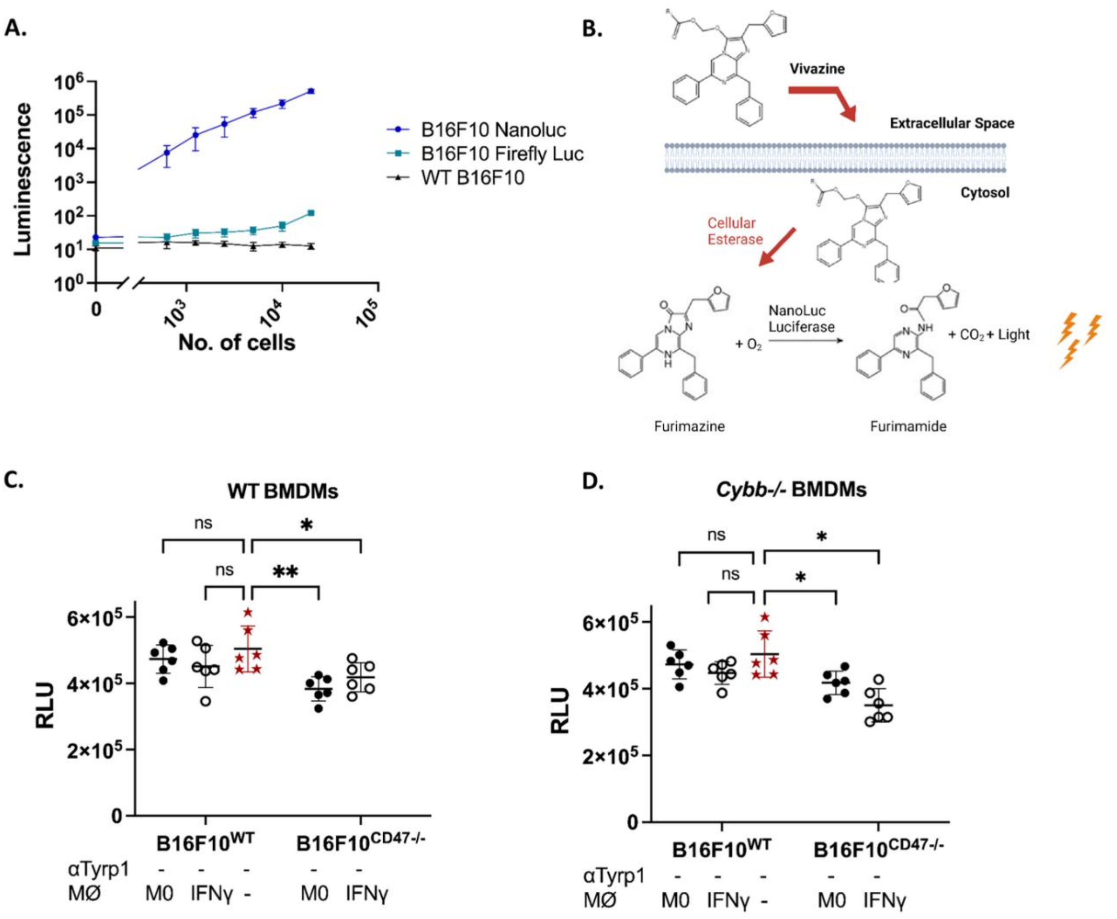


**Figure S4. Comparison of Nanoluciferase and firefly luciferase sensitivity and validation of nanoluc-based B16F10 depletion assay in WT and *Cybb*$^{-/-}$ BMDMs.**

(A) Comparison of luminescence signals generated from B16F10$^{WT}$, B16F10$^{nanoluc}$, and B16F10$^{fireflyluc}$ cells. Luminescence was measured following incubation with the substrate vivazine. Each dot represents the mean of 6 technical replicates from 2 independent experiments. (error bars ± standard deviation).

(B) Schematic illustration of NanoLuc-mediated bioluminescence. The cell-permeable substrate vivazine diffuses across the plasma membrane and is processed by the intracellular esterases to generate furimazine in the cytosol. NanoLuc catalyses the oxidation of furimazine to furimamide, resulting in light emission that can be detected as luminescence by the plate reader.

(C, D) Macrophage-mediated depletion of B16F10$^{WT}$ and B16F10$^{CD47-/-}$ tumoroids by BMDMs cultured in the absence of targeting antibody. Tumoroids were co-cultured with homeostatic M0 or IFNγ-polarized BMDMs isolated from WT (C) or *Cybb*$^{-/-}$ (D) mice. Each point represents a single tumoroid, mean ± SD from three replicates across two experiments. Statistical significance was analyzed by two-way ANOVA with Tukey's multiple comparison test. ns, non-significant; *$p < 0.05$, **$p < 0.01$.

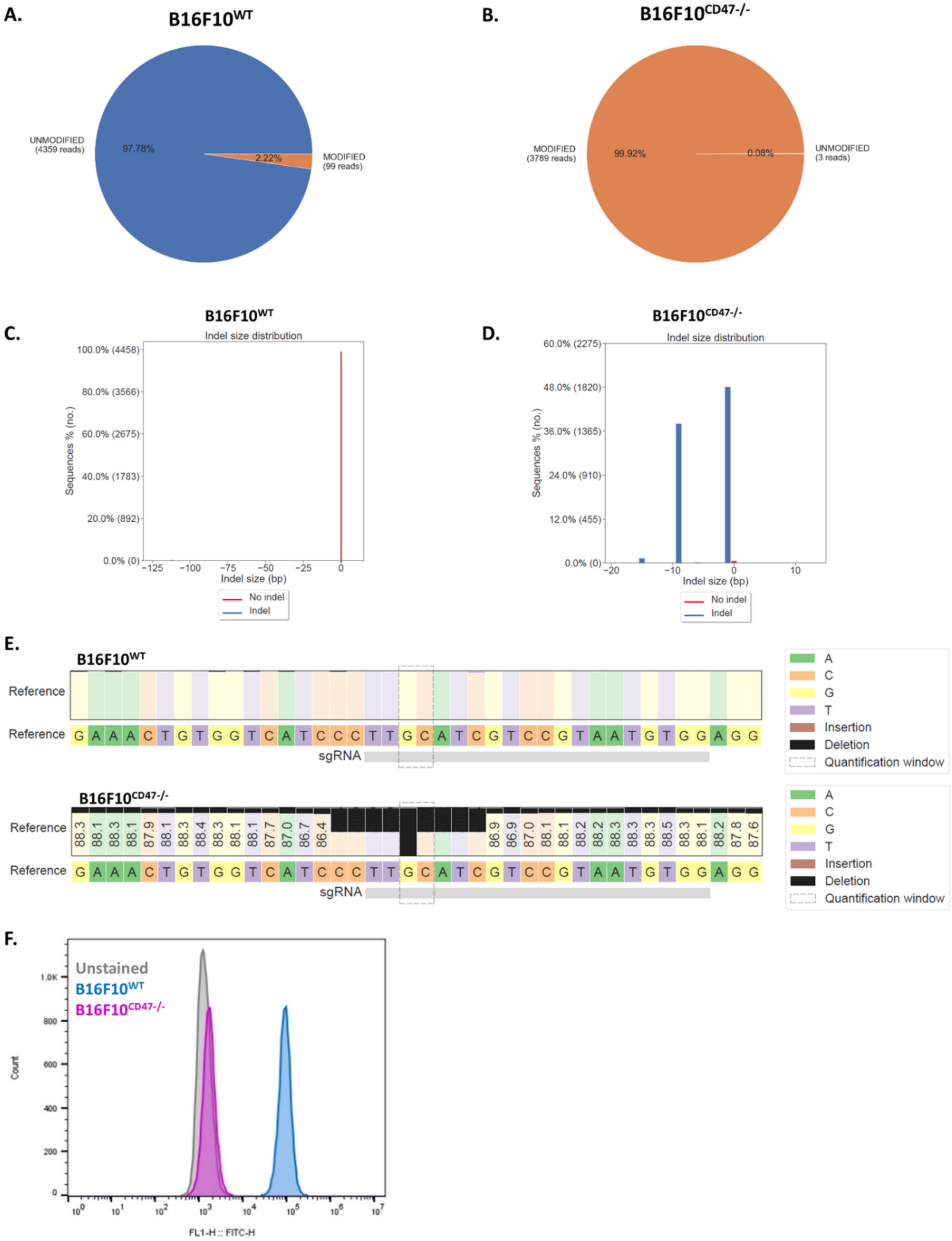

A.
B16F10WT
UNMODIFIED (4359 reads)
97.78%
2.22%
MODIFIED (99 reads)
B.
B16F10CD47-/-
MODIFIED (3789 reads)
99.92%
0.08%
UNMODIFIED (3 reads)
C.
B16F10WT
Indel size distribution
Sequences % (no.)
100.0% (4458)
80.0% (3566)
60.0% (2675)
40.0% (1783)
20.0% (892)
0.0% (0)
−125 −100 −75 −50 −25 0
Indel size (bp)
No indel
Indel
D.
B16F10CD47-/-
Indel size distribution
Sequences % (no.)
60.0% (2275)
48.0% (1820)
36.0% (1365)
24.0% (910)
12.0% (455)
0.0% (0)
−20 −10 0 10
Indel size (bp)
No indel
Indel
E.
B16F10WT
Reference
Reference G A A A C T G T G G T C A T C C C T T G C A T C G T C C G T A A T G T G G A G G
sgRNA
A
C
G
T
Insertion
Deletion
Quantification window
B16F10CD47-/-
Reference 88.3 88.1 88.3 88.1 87.9 88.1 88.3 88.4 88.3 88.1 88.1 87.7 87.0 86.7 86.4 86.9 86.9 87.0 88.1 88.1 88.2 88.2 88.3 88.3 88.3 88.5 88.3 88.1 88.2 87.8 87.6
Reference G A A A C T G T G G T C A T C C C T T G C A T C G T C C G T A A T G T G G A G G
sgRNA
A
C
G
T
Insertion
Deletion
Quantification window
F.
Unstained
B16F10WT
B16F10CD47-/-
Count
1.0K
800
600
400
200
0
FL1-H :: FITC-H

**Figure S5. Characterization of B16F10$^{CD47-/-}$ clone.**

(A, B) Alignment pie charts comparing B16F10$^{WT}$ (A) and B16F10$^{CD47-/-}$ clone (B) sequencing results analyzed using Crispresso. Pie charts represent the proportion of reads corresponding to unmodified and modified alleles in WT and clone samples.

(C, D) Indel size distribution for B16F10$^{WT}$ (C) with no detectable insertions or deletions, and B16F10$^{CD47-/-}$ clone (D) showing predominant 1-bp and 9-bp deletions at the CRISPR target site, resulting in disruption of the coding sequence and alteration of the protein structure.

(E) Nucleotide percentage quilt around sgRNA C CAC ATT ACG GAC GAT GCA A, for B16F10$^{WT}$ and B16F10$^{CD47-/-}$ clone. The WT sequence is unaltered, while the gene-edited clone exhibited deletions of nucleotides corresponding to three amino acids: proline, cysteine, and isoleucine.

(F) CD47 surface staining of B16F10$^{WT}$ and B16F10$^{CD47-/-}$ tumor cells. Cells were stained with anti-CD47 MIAP301 antibody followed by DyLight 488-conjugated goat anti-rat IgG secondary antibody. Unstained B16F10$^{WT}$ cells (grey), stained B16F10$^{WT}$ cells (blue) and B16F10$^{CD47-/-}$ cells (magenta).

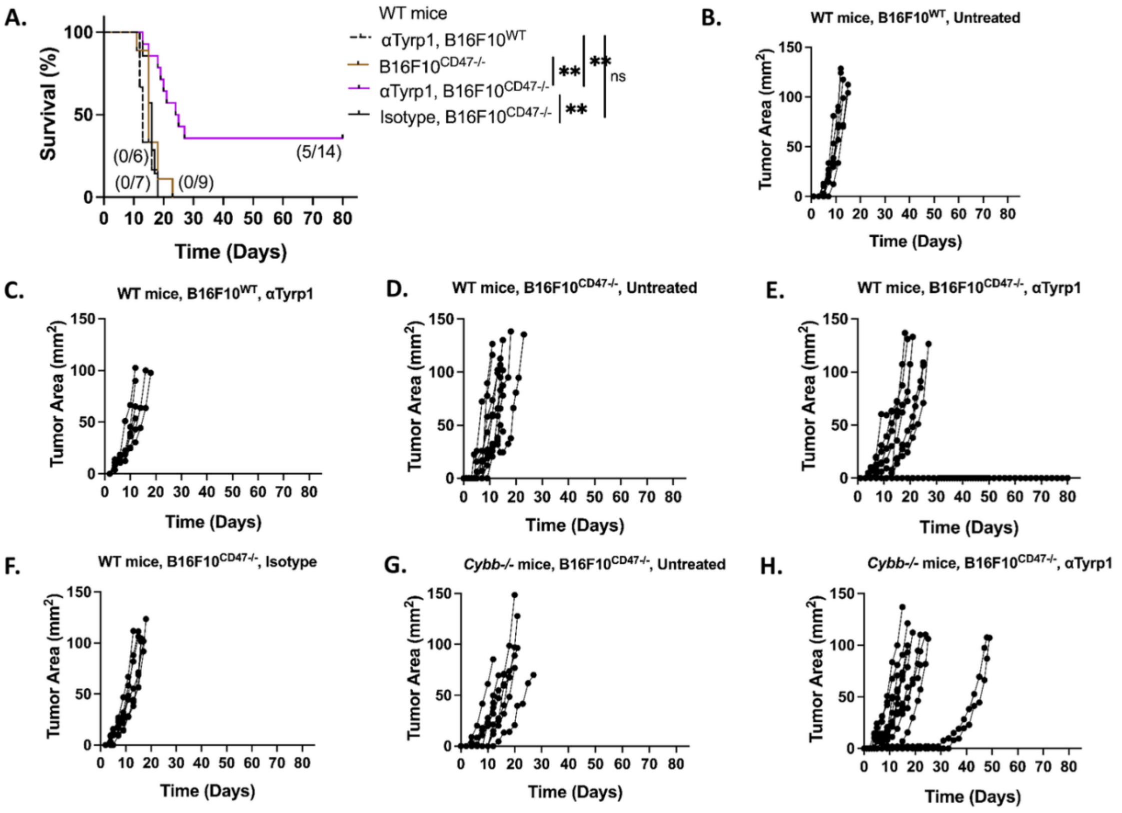


**Figure S6. Survival and tumor growth curves of WT and Cybb$^{-/-}$ mice bearing B16F10$^{WT}$ or B16F10$^{CD47-/-}$ tumors treated with anti-Tyrp1, an isotype control, or untreated.**

(A) Survival curves of WT mice bearing B16F10$^{CD47-/-}$ tumors that were left untreated (brown), treated with isotype control antibody (black), or treated with anti-Tyrp1 antibody (magenta). Survival curve of anti-Tyrp1-treated WT mice bearing B16F10$^{WT}$ tumors is shown for comparison (black dotted). Fractions in parentheses indicate the number of surviving mice over the total number of mice in each cohort. Statistical significance was determined by the Log rank (Mantel-Cox) test. ns, non-significant; $^{*}p < 0.05$, $^{**}p < 0.01$.

(B-J) Individual tumor growth curves during primary tumor challenge. WT mice bearing B16F10$^{WT}$ tumors were left untreated (B, n= 7) or treated with anti-Tyrp1 (C, n=6). WT mice bearing B16F10$^{CD47-/-}$ tumors were left untreated (D, n= 9), treated with anti-Tyrp1 (E, n=14), or treated with isotype control antibody (F, n=7). Cybb$^{-/-}$ mice bearing B16F10$^{CD47-/-}$ tumors were left untreated (G, n= 9) or treated with anti-Tyrp1 (H, n=13). Each line represents an individual mouse.

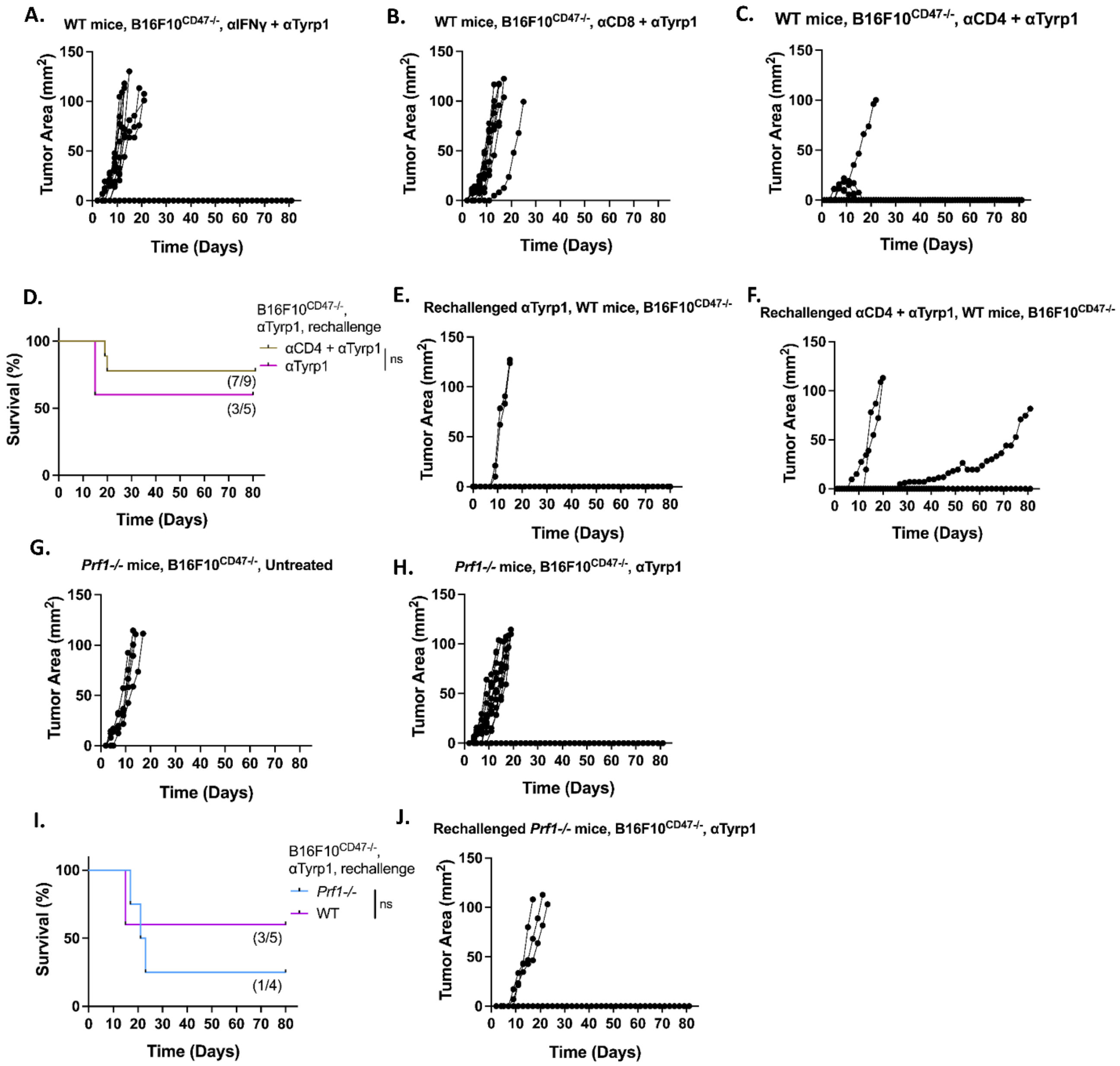

A.
WT mice, B16F10CD47-/-, αIFNγ + αTyrp1
B.
WT mice, B16F10CD47-/-, αCD8 + αTyrp1
C.
WT mice, B16F10CD47-/-, αCD4 + αTyrp1
Tumor Area (mm2)
Time (Days)
D.
B16F10CD47-/-, αTyrp1, rechallenge
αCD4 + αTyrp1
αTyrp1
ns
(7/9)
(3/5)
Survival (%)
E.
Rechallenged αTyrp1, WT mice, B16F10CD47-/-
F.
Rechallenged αCD4 + αTyrp1, WT mice, B16F10CD47-/-
G.
Prf1-/- mice, B16F10CD47-/-, Untreated
H.
Prf1-/- mice, B16F10CD47-/-, αTyrp1
I.
B16F10CD47-/-, αTyrp1, rechallenge
Prf1-/-
WT
ns
(3/5)
(1/4)
J.
Rechallenged Prf1-/- mice, B16F10CD47-/-, αTyrp1

**Figure S7. Tumor growth curves of WT mice following CD4, CD8, or IFNγ depletion, and *Cybb*$^{-/-}$ and *Prf1*$^{-/-}$ mice treated with anti-Tyrp1 and subsequent untreated rechallenge.**

(A-C) Individual tumor growth curves during primary challenge in WT mice bearing B16F10$^{CD47-/-}$ tumors treated with anti-Tyrp1 in combination with anti-IFNγ (A, n= 10), anti-CD8 (B, n= 10), or anti-CD4 (C, n=10). Each line represents an individual mouse.

(D) Survival analysis following rechallenge with B16F10$^{CD47-/-}$ cells in the absence of further treatment. Mice were survivors of the primary challenge and had previously received either anti-Tyrp1 monotherapy (magenta) or anti-Tyrp1 combined with CD4 depletion (olive). Fractions in parentheses indicate the number of surviving mice over the total number of mice in each cohort. Statistical significance was determined by the Log rank (Mantel-Cox) test.

(E, F) Individual tumor growth curves during rechallenge of WT mice that survived the primary challenge. Mice were re-inoculated with B16F10$^{CD47-/-}$ cells and received no additional therapy. Mice were previously treated with either anti-Tyrp1 initial monotherapy (E, n=5) or anti-Tyrp1 and anti-CD4 combination therapy (F, n=9).

(G, H) Individual tumor growth curves during primary challenge in *Prf1*$^{-/-}$ mice, bearing B16F10$^{CD47-/-}$ tumors that were left untreated (G, n= 5) or treated with anti-Tyrp1 (H, n= 16).

(I) Survival analysis following tumor rechallenge with B16F10$^{CD47-/-}$ cells in the absence of further treatment. *Prf1*$^{-/-}$ mice were survivors of the primary challenge after anti-Tyrp1 monotherapy (blue).

(J) Individual tumor growth curves during re-challenge of *Prf1*$^{-/-}$ mice that survived the primary challenge after anti Tyrp1 monotherapy (J, n= 4). Mice were re-inoculated with B16F10$^{CD47-/-}$ cells and received no additional therapy. ns, non-significant; *$p < 0.05$, **$p < 0.01$.

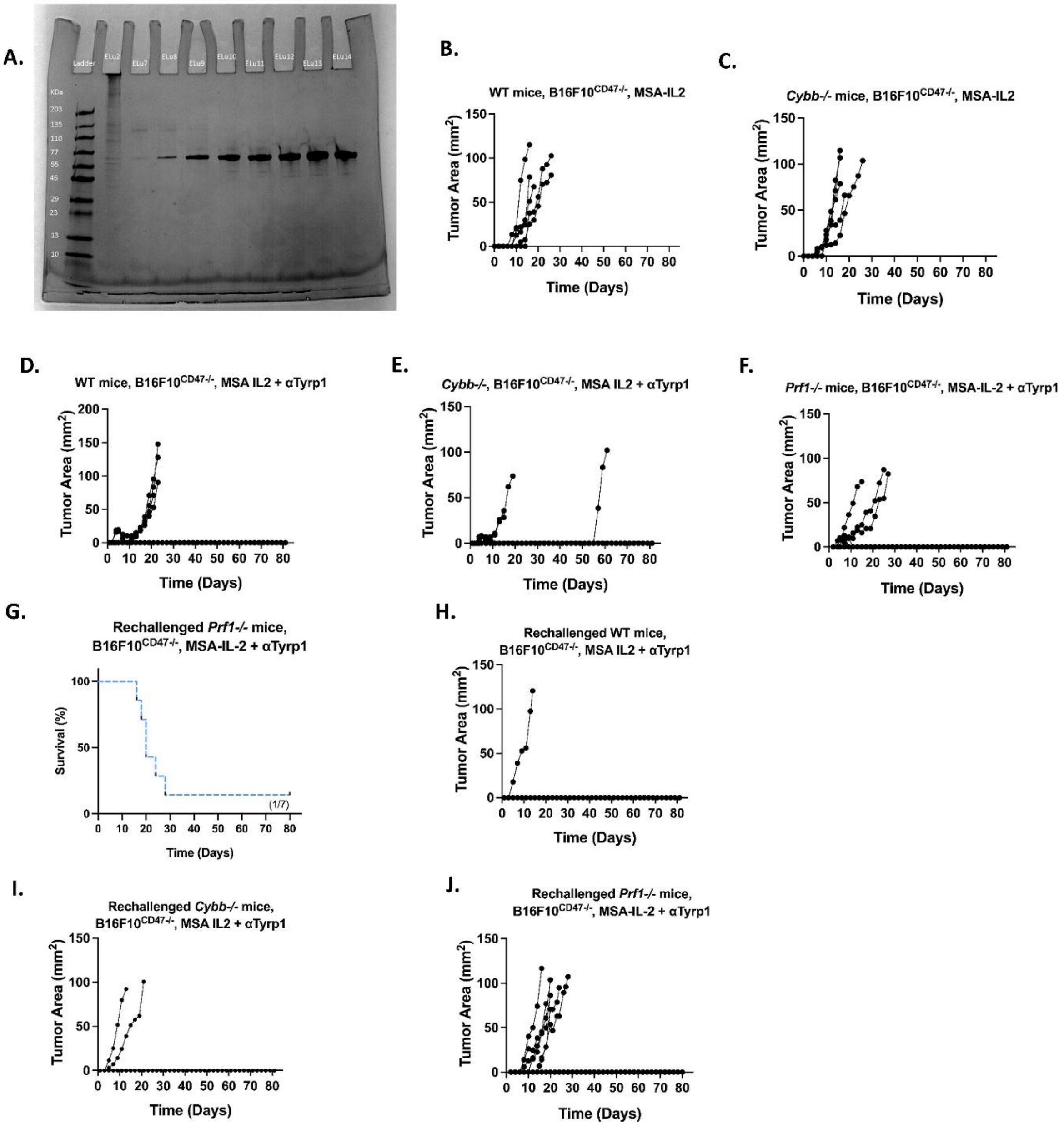
A.
Ladder
ELu2
ELu7
ELu8
ELu9
ELu10
ELu11
ELu12
ELu13
ELu14
KDa
203
135
110
77
55
46
29
23
13
10
B.
WT mice, B16F10$^{CD47-/-}$, MSA-IL2
Tumor Area (mm$^2$)
Time (Days)
C.
*Cybb-/-* mice, B16F10$^{CD47-/-}$, MSA-IL2
Tumor Area (mm$^2$)
Time (Days)
D.
WT mice, B16F10$^{CD47-/-}$, MSA IL2 + αTyrp1
Tumor Area (mm$^2$)
Time (Days)
E.
*Cybb-/-*, B16F10$^{CD47-/-}$, MSA IL2 + αTyrp1
Tumor Area (mm$^2$)
Time (Days)
F.
*Prf1-/-* mice, B16F10$^{CD47-/-}$, MSA-IL-2 + αTyrp1
Tumor Area (mm$^2$)
Time (Days)
G.
Rechallenged *Prf1-/-* mice,
B16F10$^{CD47-/-}$, MSA-IL-2 + αTyrp1
Survival (%)
(1/7)
Time (Days)
H.
Rechallenged WT mice,
B16F10$^{CD47-/-}$, MSA IL2 + αTyrp1
Tumor Area (mm$^2$)
Time (Days)
I.
Rechallenged *Cybb-/-* mice,
B16F10$^{CD47-/-}$, MSA IL2 + αTyrp1
Tumor Area (mm$^2$)
Time (Days)
J.
Rechallenged *Prf1-/-* mice,
B16F10$^{CD47-/-}$, MSA-IL-2 + αTyrp1
Tumor Area (mm$^2$)
Time (Days)

**Figure S8. MSA-IL-2 purification and tumor growth curves of WT, *Cybb*$^{-/-}$, and *Prf1*$^{-/-}$ mice treated with MSA-IL-2 alone or in combination with anti-Tyrp1 and subsequent untreated rechallenge.**

(A) SDS-PAGE analysis of purified MSA-IL-2 following size-exclusion chromatography.

(B, C) Individual tumor growth curves of WT mice (B, n=5) or *Cybb*$^{-/-}$ mice (C, n=5), bearing B16F10$^{CD47-/-}$ tumors and treated with MSA-IL2 monotherapy. Each line represents an individual mouse.

(D-F) Individual tumor growth curves of individual WT (D, n=9) or *Cybb*$^{-/-}$ (E, n=10) or *Prf1*$^{-/-}$ mice (F, n=10), bearing B16F10$^{CD47-/-}$ tumors and treated with MSA-IL2 and anti-Tyrp1 combination therapy.

(G) Survival analysis following re-challenge with B16F10$^{CD47-/-}$ tumors in the absence of further treatment. Re-challenged *Prf1*$^{-/-}$ (blue dotted) mice were survivors of the primary challenge after MSA-IL2 and anti-Tyrp1 combination therapy.

(H-J) Individual tumor growth curves during re-challenge of survivors from the primary challenge. Mice were re-inoculated with B16F10$^{CD47-/-}$ cells and received no additional therapy. WT (H, n=5) or *Cybb*$^{-/-}$ (I, n=5) or *Prf1*$^{-/-}$ (J, n=7) mice were previously treated with MSA-IL2 and anti-Tyrp1 combination therapy.

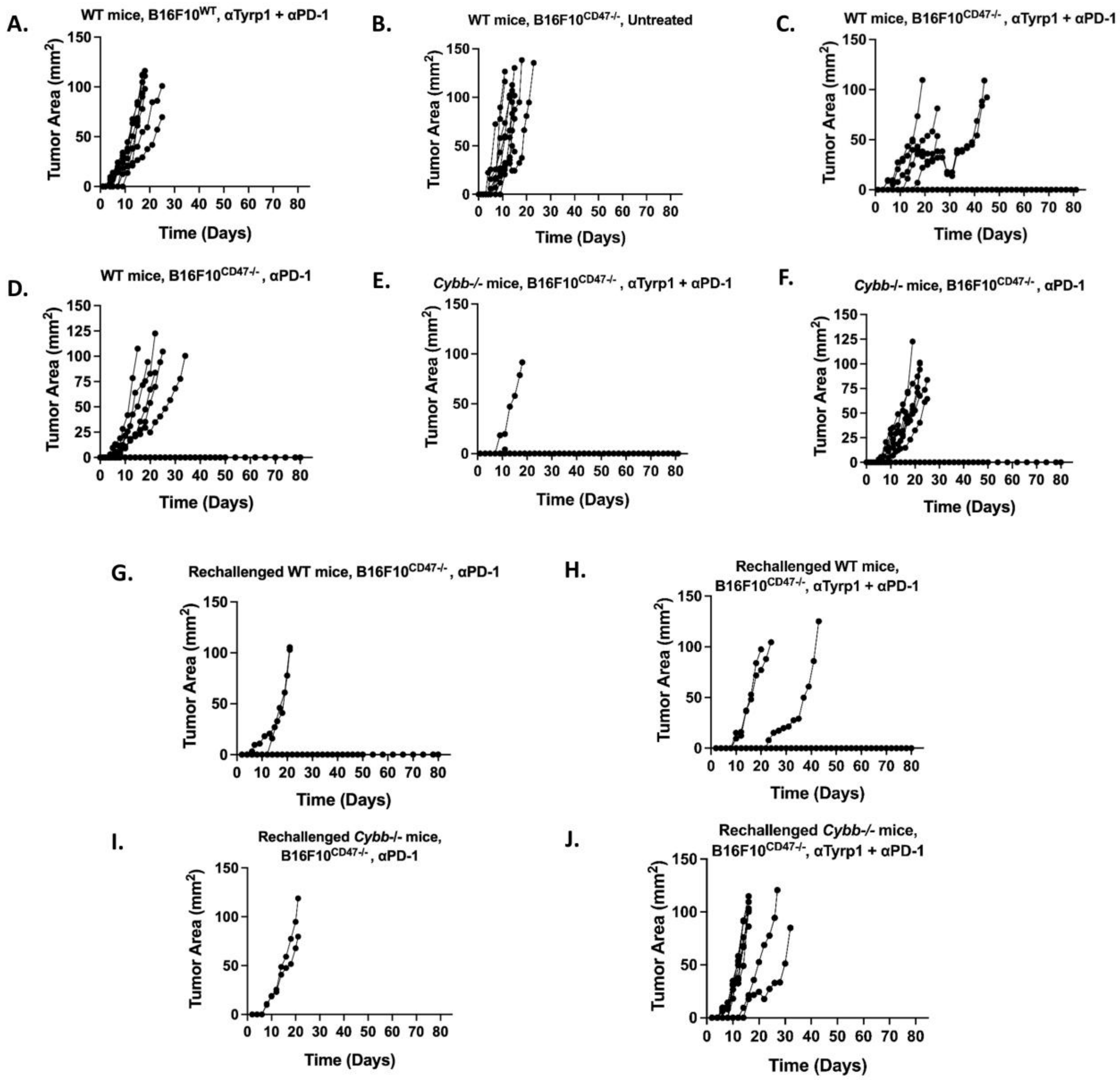
A.
WT mice, B16F10^WT, αTyrp1 + αPD-1
B.
WT mice, B16F10^CD47-/-, Untreated
C.
WT mice, B16F10^CD47-/-, αTyrp1 + αPD-1
D.
WT mice, B16F10^CD47-/-, αPD-1
E.
Cybb-/- mice, B16F10^CD47-/-, αTyrp1 + αPD-1
F.
Cybb-/- mice, B16F10^CD47-/-, αPD-1
G.
Rechallenged WT mice, B16F10^CD47-/-, αPD-1
H.
Rechallenged WT mice, B16F10^CD47-/-, αTyrp1 + αPD-1
I.
Rechallenged Cybb-/- mice, B16F10^CD47-/-, αPD-1
J.
Rechallenged Cybb-/- mice, B16F10^CD47-/-, αTyrp1 + αPD-1
Tumor Area (mm2)
Time (Days)

**Figure S9. Tumor growth curves of WT and *Cybb*$^{-/-}$ mice treated with anti-PD-1 alone or in combination with anti-Tyrp1 and subsequent untreated rechallenge.**

(A-H) Individual tumor growth curves during primary challenge. WT mice bearing B16F10$^{WT}$ tumors were treated with anti-PD-1 and anti-Tyrp1 combined (A, n=8) or anti-PD-1 alone (B, n= 5). WT mice bearing B16F10$^{CD47-/-}$ tumors were treated with anti-PD-1 and anti-Tyrp1 combination therapy (C, n=10) or anti-PD-1 monotherapy (D, n= 10). *Cybb*$^{-/-}$ mice bearing B16F10$^{CD47-/-}$ tumors were treated with anti-PD-1 and anti-Tyrp1 combined (E, n=10) or anti-PD-1 alone (F, n= 10). Each line represents an individual mouse.

(G-J) Individual tumor growth curves following tumor re-challenge with B16F10$^{CD47-/-}$ cells in the absence of further treatment. Mice were survivors from the primary challenge. WT cohort consisted of survivors previously treated with either anti-PD-1 alone (G, n=4) or anti-PD-1 and anti-Tyrp1 combined (H, n=5). *Cybb*$^{-/-}$ cohort consisted of survivors previously treated with either anti-PD-1 alone (I, n=2) or anti-PD-1 and anti-Tyrp1 combined (J, n=9).

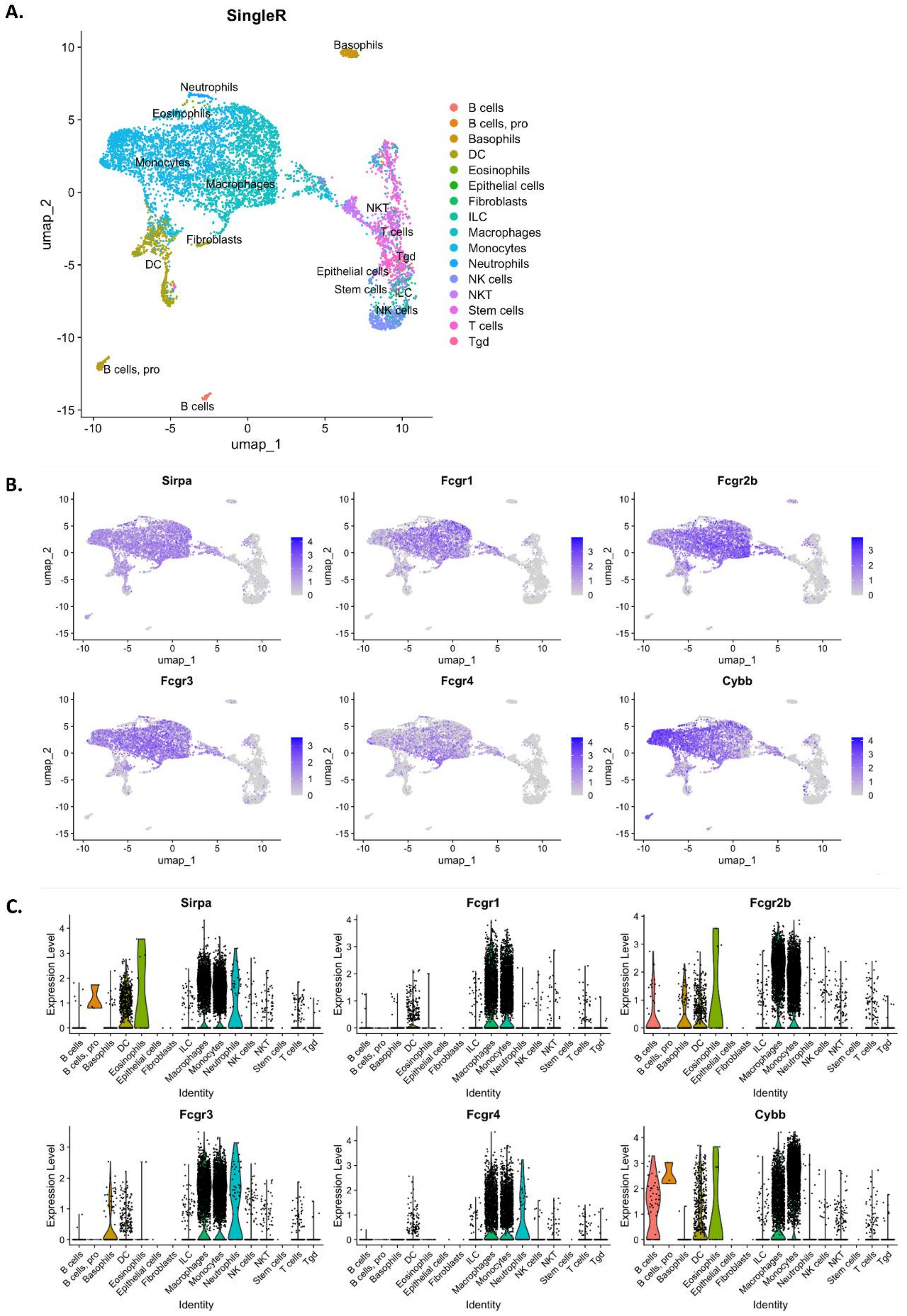
A.
SingleR
B cells
B cells, pro
Basophils
DC
Eosinophils
Epithelial cells
Fibroblasts
ILC
Macrophages
Monocytes
Neutrophils
NK cells
NKT
Stem cells
T cells
Tgd
umap_1
umap_2
B.
Sirpa
Fcgr1
Fcgr2b
Fcgr3
Fcgr4
Cybb
C.
Expression Level
Identity

**Figure S10. Expression of ADCP-related genes in immune populations of B16F10 tumors.**

(A-C) CD45-positive immune cell abundance, *Sirpa*, FcγRs, and *Cybb* expression were quantified in late-stage B16F10 (day 16, untreated) melanoma tumors by single-cell RNA-seq (GSM8479167, Yuanhanyu Luo 2024) using Seurat v5 (Satija Lab), and SingleR (Immgen).
